# Age and sex dependent variability of type 2 dopamine receptors in the human brain: A large-scale PET cohort

**DOI:** 10.1101/2021.08.10.455776

**Authors:** Tuulia Malén, Tomi Karjalainen, Janne Isojärvi, Aki Vehtari, Paul-Christian Bürkner, Vesa Putkinen, Valtteri Kaasinen, Jarmo Hietala, Pirjo Nuutila, Juha Rinne, Lauri Nummenmaa

**Author notes:** **Address Correspondence to:** Tuulia Malén, Turku PET Centre c/o Turku University Hospital, FI-20520 Turku, Finland.

## Abstract

**BACKGROUND:** The dopamine system contributes to a multitude of functions ranging from reward and motivation to learning and movement control, making it a key component in goal-directed behavior. Altered dopaminergic function is observed in neurological and psychiatric conditions. Numerous factors have been proposed to influence dopamine function, but due to small sample sizes and heterogeneous data analysis methods in previous studies their specific and joint contributions remain unresolved.

**METHODS:** In this cross-sectional register-based study we investigated how age, sex, body mass index (BMI), as well as cerebral hemisphere and regional volume influence striatal type 2 dopamine receptor (D_2_R) availability in the human brain. We analyzed a large historical dataset (n=156, 120 males and 36 females) of [11C]raclopride PET scans performed between 2004 and 2018.

**RESULTS:** Striatal D_2_R availability decreased through age for both sexes and was higher in females versus males throughout age. BMI and striatal D_2_R availability were weakly associated. There was no consistent lateralization of striatal D_2_R. The observed effects were independent of regional volumes. These results were validated using two different spatial normalization methods, and the age and sex effects also replicated in an independent sample (n=135).

**CONCLUSIONS:** D_2_R density is dependent on age and sex, which may contribute to the vulnerability of neurological and psychiatric conditions involving altering D_2_R expression.

## Introduction

Dopaminergic function regulates emotion, cognition and learning as well as motor functions (1, 2), making the dopamine system a key component for goal-directed behavior (3–5). Aberrant dopaminergic function is observed in various neurological and psychiatric conditions, such as Parkinson’s disease, schizophrenia, drug abuse, obesity and depression (6–8). Dopamine receptors are divided into type 1 (D_1_R including types D_1_ and D_5_) and type 2 (D_2_R including types D_2_, D_3_ and D_4_) receptor families (9, 2). Particularly the D_2_R which is abundantly expressed in the striatum (10, 11) is centrally involved in the pathophysiology of neuropsychiatric conditions (7).

Patients with schizophrenia show striatal hyperactivity of dopaminergic function (12) and elevated D_2_R density (13), yet it remains unresolved how the disorder itself (e.g. illness duration) and exposure to antipsychotic medication link to these observations (14). D_2_Rs also mediate anxious symptomology (15, 16) and elevated D_2_R expression is observed in motivational disturbance (17) and possibly in depression, although as the elevated D_2_R has been shown particularly in medicated (18) rather than unmedicated depression (19), possibly reflecting antidepressant treatment (19). Conversely, Parkinson’s disease is associated with lowered D_2_R expression (7), at least after the early disorder stage of when increase of D_2_Rs may occur as a compensation to nerve terminal loss or medical treatment (20, 13). In addition to this neurodegenerative disease (7, 21), drug abuse is also associated with striatal D_2_R loss, and the lower D_2_R density may constitute a vulnerability factor for drug abuse (22, 23).

To understand dopaminergic dysfunction and related pathophysiology, factors contributing to dopamine function in the healthy population need to be identified. Small-scale PET studies suggest that subject demographics, such as age (24–27), sex (28, 29) and body mass index (BMI) (27, 30, but see 31) might affect the D_2_R availability in striatum. However, there has been increasing concern over the lack of replicability of neuroimaging findings (32). Insufficient statistical power (32, 33), variable methods for analyzing data (34), as well as failure to appropriately control for multiple comparisons (35) have been proposed as main sources of the poor replicability.

Because PET imaging is expensive, data pooling has recently emerged as an effective way of increasing sample sizes and consequently providing accurate statistical estimates (36). Additionally, Bayesian hierarchical modeling has been proposed to facilitate reproducible science by limiting the “researcher degrees of freedom” in the analysis phase (37) and by removing the need for arbitrary multiple comparison correction methods (38). Here we used Bayesian hierarchical modeling to estimate how age, sex, BMI and regional volume are associated with D_2_R cerebral lateralization. We analyzed a large dataset of 156 historical controls scanned with [11C]raclopride, a selective antagonist D_2_R radioligand. We also replicated the results in an independent sample of 135 subjects.

## Methods and Materials

### Subjects

The data were 156 baseline [11C]raclopride scans of healthy control subjects (sex 120 males and 36 females; age 19-71 years, BMI range 18-38) scanned at Turku PET Centre between 2004 and 2018. Detailed sample information is shown in **Table 1** (see also Table S3 for exclusion criteria). Studies were included in the analysis if they were baseline scans with injected dose > 100 MBq (to avoid low signal-noise ratio (SNR)) and the magnetic resonance (MR) scan and basic demographic and anthropometric information (height, weight) was available. If multiple baseline scans were acquired for an individual, chronologically first scan was included in the analysis. Finnish legislation does not require ethical approval for register-based studies.

**Table 1.** Characteristics of the sample. See Supplementary Material (SM) section Scanner Considerations for scanner information and Table S3 for exclusion criteria. SD= standard deviation. Note that using HRRT, the injected activity is significantly higher compared with the other scanners, explaining the high SD in injected activity.

|  | Males (n= 120) |  |  | Females (n= 36) |  |  |
| --- | --- | --- | --- | --- | --- | --- |
|  | Mean | SD | Range | Mean | SD | Range |
| Injected Activity (MBq) | 325 | 118 | 183-538 | 258 | 39 | 187-329 |
| Age (years) | 25 | 5 | 19-56 | 37 | 14 | 20-71 |
| Height (cm) | 181 | 7 | 167-199 | 166 | 7 | 151-190 |
| Weight (kg) | 79 | 12 | 58-130 | 61 | 8 | 47-85 |
| BMI (kg/m<sup>2</sup>) | 24 | 3 | 19-38 | 22 | 3 | 18-33 |

### PET data acquisition and Image Processing

Antagonist radioligand [11C]raclopride binds to D_2_Rs (39–41), allowing reliable quantification of striatal and thalamic D_2_R availability (42–45, 41, 46). However, its binding in extrastriatal regions, such as the cerebral cortex, is unspecific (44, 47, but see 46, 48). In this study we included the following four regions of interest (ROIs): striatal nucleus accumbens (accumbens), caudate nucleus (caudate), putamen, as well as thalamus close by the striatal ROIs. The PET data was acquired using five different scanners (Scanner Considerations in SM).

Preprocessing and kinetic modeling were done using Magia toolbox (49). Preprocessing consisted of framewise realignment and co-registration of the PET and magnetic resonance images (MRIs). Tracer binding was quantified using the outcome measure binding potential (BP_ND_), which is the ratio of specific binding to non-displaceable binding in tissue (45). BP_ND_ was estimated using a simplified reference tissue model (50) with cerebellar gray matter as the reference region (51). Data length was harmonized by including first 52 minutes from each scan. Individual frames were first realigned to account for between-frame movements. The first frame was because it did not contain sufficient signal for every subject. T1-weighted MR images were processed using FreeSurfer (https://surfer.nmr.mgh.harvard.edu/). The MR images were then co-registered with the PET data for region of interest extraction.

### Statistical modeling

Statistical modeling was carried out in R (52) using brms (53, 54) that applies the Markov-Chain Monte Carlo sampling tools of RStan (55). The analysis script is available in SM code.

#### Primary analysis

We first standardized the continuous variables and log-transformed binding potential estimates because according to posterior predictive checks (56, 57), log-transformation of non-negative dependent variable enhances model fit, as it makes the model multiplicative instead of additive that is not optimal when limited to positive values (58). We also confirmed that the age and BMI effects on logarithmic BP_ND_ are well approximated by a linear function in each ROI (Linearity Assessment of the Age and BMI effects in SM). For the sake of conciseness, we simply refer to the linear effects on a logarithmic scale as linear. We used Bayesian hierarchical regression to model the data. Because ROI-wise effects were partially pooled across ROIs, this essentially removes the need to correct for multiple comparisons induced by investigating multiple ROIs (59).

We estimated one primary model for assessing the main effects of age, sex and BMI on BP_ND_. The effects were calculated separately for the left and right hemisphere. We also investigated the main effect of cerebral hemisphere (i.e. lateralization) on BP_ND_ separately for males and females, as our initial modeling showed sex-differences in lateralization and as previous research has pointed to greater lateralization in the male versus female brain (60–62). There was no clear evidence for sex-specific age-effect (Figure S6), prompting us to calculate the age effect together for males and females with maximal statistical power. To estimate the effects of age, sex, BMI and hemisphere, we used regionally varying random slopes. Subjects, scanners, and ROIs, were all modelled as varying (random) intercepts. We included a varying intercept for the combination of scanners and ROIs, to allow for regionally varying scanner effects (Scanner Considerations in SM). For the residual variances, we applied the same grouping structure, except for subject (no individual differences expected). Additional modeling information is presented in SM (Sampling Settings and Convergence Estimates).

#### Sensitivity analysis and replication in an independent sample

The large number of young male subjects resulted in an imbalanced sex ratio, especially after the age of 40. Hence, we repeated the primary model with a balanced subset of the data (n= 140, see Tables S1 and S3), including the data of subjects aged 40 and under. We also checked whether adjusting for inter-individual differences in regional volumes of the ROIs changed the results or had main effects on D_2_R binding (See SM file).

Our independent secondary sample of 135 scans (104 males, mean age 33 years, Tables S2 and S3) was not included in the primary analysis due to missing MR images or anthropometric measurements (weight, height) from these subjects. In a secondary analysis we maximized statistical power by applying template-based normalization method to the whole available sample (primary and secondary). We first validated the normalization and ROI extraction protocol without the MR images by conducting a within-subject comparison of the BP_ND_ estimates produced by the two normalization methods for the subjects that both normalization methods could be applied (MR image available, n= 189, Table S3). The analysis showed that both methods yield comparable BP_ND_ estimates (Pearson’s product-moment correlation coefficients 0.97-0.99). Then, we replicated the statistical analysis of the global effects of age and sex on the D_2_R BP_ND_ (using template-based normalization method) using the secondary sample with no available MRI image. See SM for more detailed information about the samples (Tables S2 and S3), validation and replication (Validation of an alternative approach for defining ROIs and reference regions).

## Results

The[11C]raclopride binding was highest in striatum and practically nonexistent in the cortex (Figure 1). The BP_ND_ in the ROIs varies from below 1 to above 5, being lowest in thalamus and highest in putamen (Figure 2). Please see **Figure S12** for the correlation of the BP_ND_ estimates between the ROIs.

**Figure 1.**
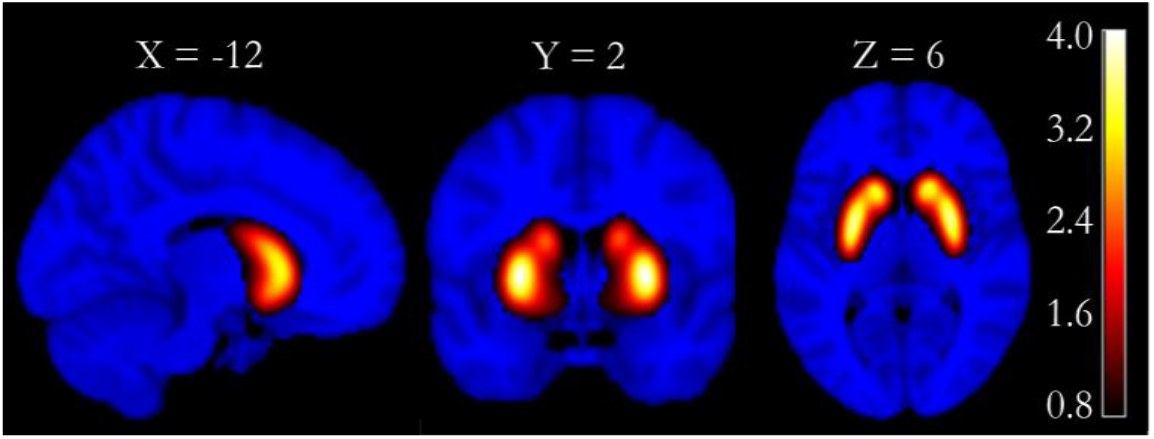
Mean [11C]raclopride BP_ND_ (original scale) in the primary sample (n= 156), on an MNI template.

**Figure 2.**
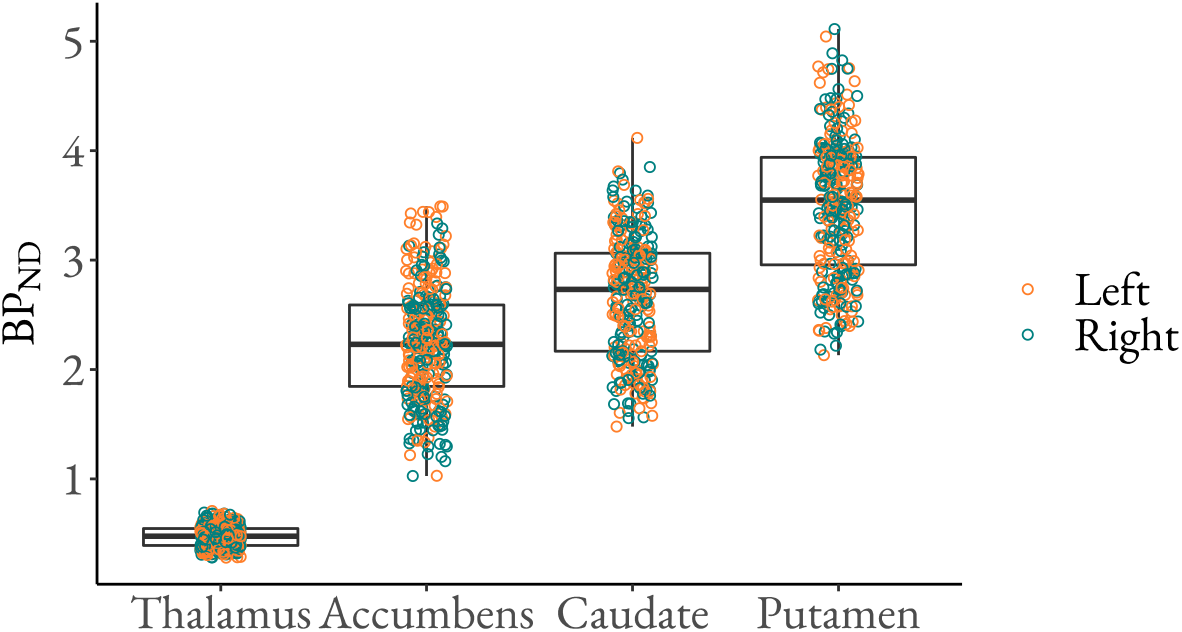
Regional [11C]raclopride BP_ND_ (original scale). The figure shows medians (middle line), 25% (lower hinge) and 75% (upper hinge) quantiles, min value (lower whisker) and max value (upper whisker), as well as the data points for the D_2_R BP_ND_.

### Age and Sex

There was a consistent age-related decline in striatal D_2_R binding (Figure 3–4). This applied particularly to the age-range from early 20s to 60s for which we had sufficient data. In putamen and caudate, 10 years of ageing (one SD) decreased the binding approximately 5%. In accumbens, the approximate decrease was 2-3% per SD. Only in thalamus, the 95% posterior uncertainty interval overlapped with zero, suggesting uncertainty in the effects. These effects were similar in both hemispheres. The further assessment supported the linearity of the age effect (Linearity Assessment of the Age and BMI effects in SM) and that the effect remains clear even when adjusting for regional volumes (Figure S7).

**Figure 3.**
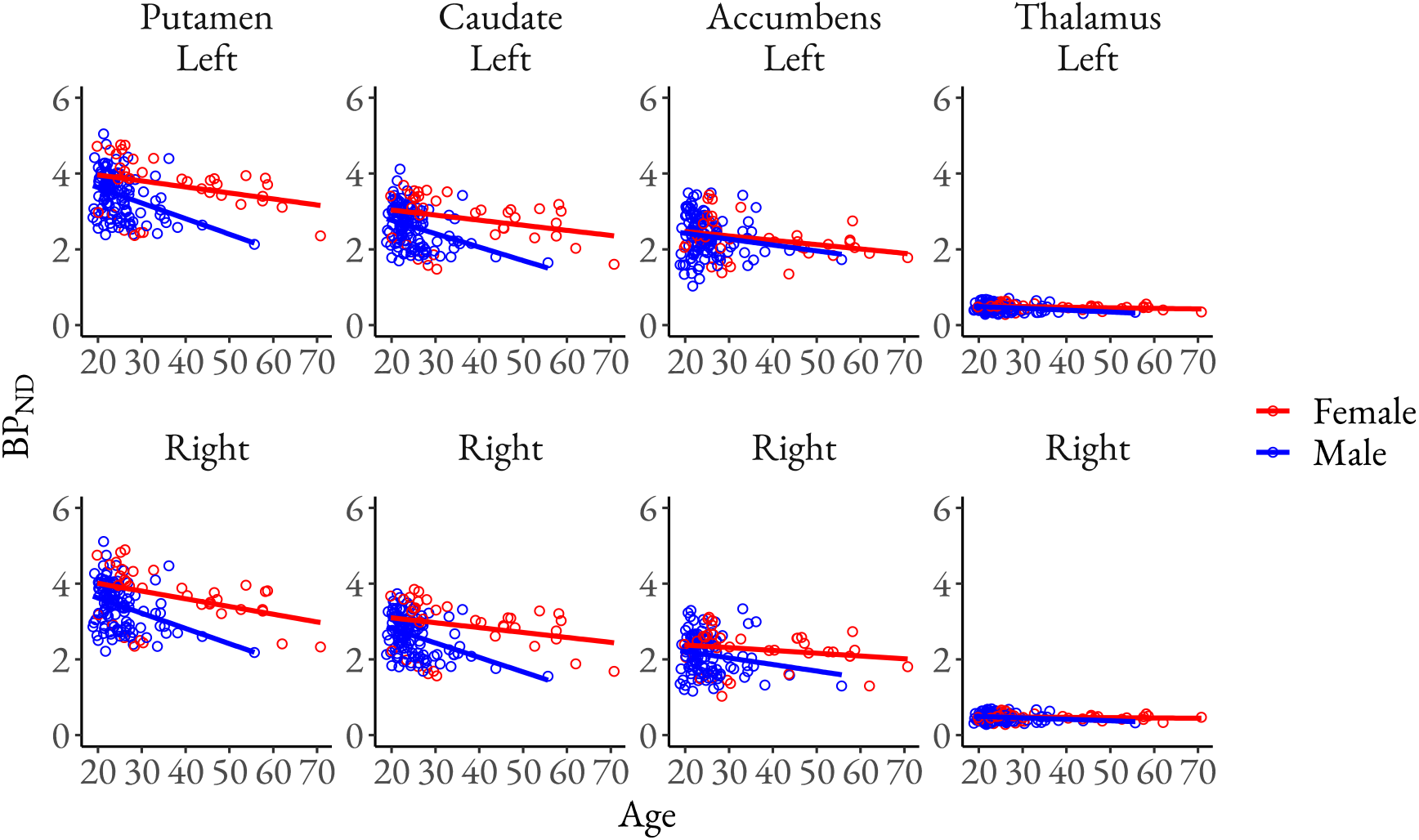
Left and right D_2_R BP_ND_ (original scale) as a function of age (original scale) in each ROI. The figure shows the original BP_ND_ estimates (points), linear regression lines separately for males and females (lines) and their 95% confidence intervals (shaded areas).

**Figure 4.**
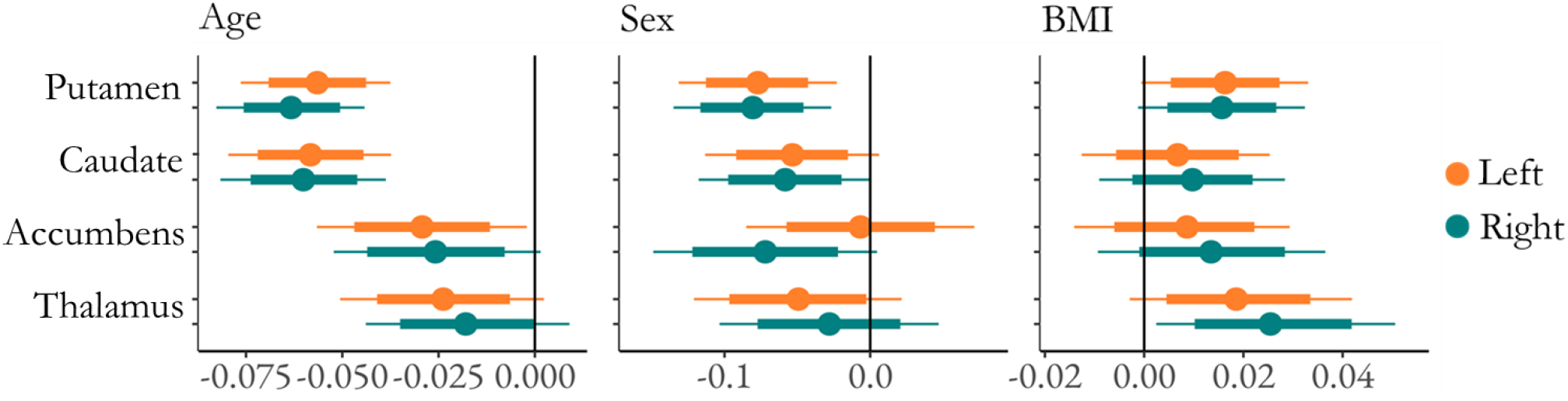
The effects of age (standardized), sex (male - female) and BMI (standardized) on striatal and thalamic D_2_R BP_ND_ (logarithmic) separately for left and right hemisphere. The figure shows medians (circles), 80% (thick line) and 95% (thin line) posterior uncertainty intervals of the regression coefficients on a logarithmic scale.

The data did not support an interactive effect of age and sex on D_2_R binding (Figure S6), instead suggesting that the age-related decline is similar for both sexes. However, the data revealed that females had on average approximately 7-8% higher D_2_R binding than males bilaterally in putamen (Figure 4). BP_ND_ tended to be higher in females versus males also in the other ROIs, although the 95% posterior uncertainty intervals overlapped with zero.

The effect of sex was in general similar in both hemispheres. Only in accumbens, the posterior uncertainty interval means for left and right hemisphere clearly differed (Figure 4), suggesting that females have higher binding than males in the right but not left hemisphere. However, the difference between the means was abolished after adjusting for regional volumes (Figure S7) and the posterior uncertainty intervals were wide in both conditions (Figures 4 and S7), reflecting uncertainty in the effects. As there was overlap in the posterior uncertainty intervals, these results do not clearly support lateralization of the sex effect even in accumbens. Adjusting for regional volumes did not change the overall effects of sex. After the adjustment for regional volumes, however, the 95% posterior uncertainty interval of right accumbens and right caudate no longer overlapped with zero, suggesting that the difference between males and females was more profound (8% in accumbens and 6% in caudate) when adjusting for regional volumes (Figure S7).

### Body Mass Index

We found no clear evidence for the effect of BMI in the D_2_R availability. However, the weak effect suggested an increase in BP_ND_ as a function of BMI across the whole range (18–38) (Figure 4). In the right thalamus, the posterior 95% uncertainty interval did not overlap with zero with an estimation of an approximate 3% increase in D_2_R binding for the increase of one SD (3 units) in BMI. In other ROIs, particularly in putamen, the majority of the posterior uncertainty intervals were above zero, also supporting the positive effect. Further assessment supported the linearity of the effects (Linearity Assessment of the Age and BMI effects in SM). Adjusting for regional volumes did not change the overall results of BMI (Figure S7).

### Lateralization

For males the data supported lateralization in all ROIs, although the direction was not coherent between the closely located ROIs (modeling results in Figure 5). The lateralization effect was strongest in accumbens, where the posterior mean and the relatively narrow posterior uncertainty interval clearly parted from zero. The binding was approximately 9% higher in left versus right accumbens. The binding was also increased in left versus right putamen, and right versus left thalamus and caudate. These effects were however smaller and the posterior uncertainty intervals overlapped zero. In females, no clear lateralization effects were found (Figure 5). The uncertainty intervals for females were wider than for males, as we had less data for females than males. Although the posterior uncertainty interval overlapped with zero, there was some support for higher binding in right versus left caudate, in line with the data from male subjects.

**Figure 5.**
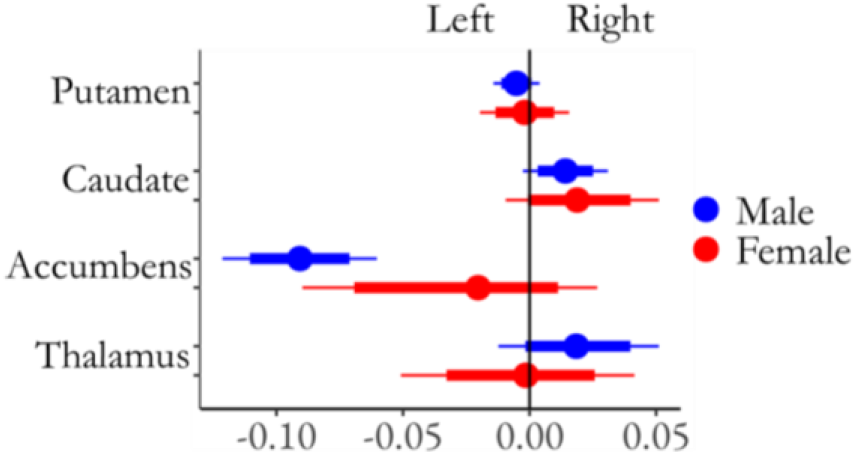
The effect of hemisphere (right - left) on striatal and thalamic D_2_R BP_ND_ (logarithmic) separately for males and females. The figure shows medians (circles), 80% (thick line) and 95% (thin line) posterior uncertainty intervals of the regression coefficients on a logarithmic scale.

### Replication analysis

The global ROI-level effects of age and sex replicated in the secondary sample (see Tables S2 and S3) that was spatially normalized with an alternative method that does not require MR image. BMI could not be included in the analysis due to missing anthropometric measurements (weight, height) for some of the subjects.

## Discussion

Our main findings were that i) there is a steady decline in D_2_R availability as a function of ageing and ii) females have higher D_2_R availability than men irrespective of age (at least from 20 to 60 years of age for which we had sufficient data). Additionally, higher BMI might be associated with increased D_2_R availability, as we found a weak positive effect of BMI. The effects of lateralization did not show clear consistence. Adjusting for volumes of the ROIs did not change the overall results, suggesting that among healthy adults, the striatal effects of age and sex on the D_2_R availability are global and independent from the regional volumes.

### Effects of Age & Sex

Our data showed a clear age-dependent decline in the striatal BP_ND_, suggesting decrease through age in D_2_Rs, starting from early adulthood. Compared to previous smaller studies (24, 25, 28, 26), our large-scale sample allowed a reliable assessment of this effect across a wide age range. Dopamine receptor loss starts already in early twenties and continues steadily throughout ageing, while previously the decline has suggested to slow down with age (24, 25, 63). The observed receptor decline changes the properties of dopamine neurotransmission (64), of which disturbance relates to several cognitive and motor symptoms (7). The etiology of Parkinson’s disease differs from mere age-related neurodegeneration (65) and doesn’t appear as accelerated aging of the dopaminergic function (66). The decline in dopamine neurotransmitter (65, 67), receptors and transporters (DATs) (68, 69, 66, 70), emerging through age, could however contribute to both the mild cognitive decline and motor deficiency commonly observed among the elderly (71, 72), as well as the more severe forms of neurodegeneration, such as Parkinsonism (7).

Our data revealed that the D_2_R availability declines similarly in both sexes, but that the average D_2_R level remains higher in females. This contrasts with prior studies that have reported sex-dependent decline in dopamine function (73), with males showing steeper reduction in receptors (28) and presynaptic dopamine synthesis (74). Our data shows that although the decline in the available D_2_Rs is not sex dependent, the overall D_2_R level is, consistently higher in females from early adulthood to at least the age 60. Sex differences have previously been observed in D_2_R affinity (lower in women) but not density, pointing to higher dopamine concentration in women (28). One study with [18F]Fluorodopa also showed greater striatal presynaptic dopamine synthesis capacity in females versus males (74).

The sex differences in the dopamine system may contribute to vulnerability for neuropsychiatric disorders (28, 74). Females might be predisposed to pathology associated with elevated D_2_R availability, such as mood disorders (75, but also consider 19). Conversely, lower D_2_R level of males may predispose them to Parkinson’s disease (75) that involves receptor loss (20, 7) and is approximately 1.5 more common in males versus females (21). Substance use disorders (76) and gambling problems (77) are significantly more prevalent in males. As conditions such as alcoholism (78) and drug abuse (23) are linked with lowered D_2_R availability, lowered D_2_R may be a potential sex-specific vulnerability factor for addictions. Finally, low striatal D_2_R density has also been associated with A1 allele of the D_2_R gene (79) that possibly links to alcoholism (80). As the deficiency in dopaminergic function does not only increase the impulsive behavior toward the object of addiction but also disturbs the saliency attribution of other objects (23), altered dopaminergic function may well constitute a significant vulnerability endophenotype for addictive behaviors. Finally, sex-differences in the dopamine system have been observed not only in the striatal (28) and cortical (81) D_2_Rs but also in DATs (66, 82) and presynaptic dopamine synthesis capacity (74). In addition, sex-specific hormones and genes play a role in the dopaminergic function and neuropsychiatric well-being (75, 67). Hence, the sex-differences in the D_2_R level may reflect broader dopaminergic, as well as dopamine related hormonal and genetic differences between sexes, and those differences may together contribute sex-dependent prevalence of neuropsychiatric disorders.

### Effect of BMI

BMI was only weakly associated with higher D_2_R availability, mainly in putamen and thalamus. Although the modeling showed uncertainty in the BMI effect, the effect was systematically positive in each ROI. As most subjects had BMI in the range of 18-30, the effects are uncertain beyond this point, thus being uninformative regarding the most seriously obese phenotypes. Previous in vivo imaging studies have yielded mixed results on the association between BMI and dopamine system (83, 27, 30, 84). Previously, decreased dopamine function (84), TaqA1 variant of D_2_R gene (85–87) and diminished incentive to physical exercise (high energy expenditure) (88) has been linked to obesity. As dopamine contributes to food-related hedonia (89), the decreased dopaminergic function could limit the rewarding effect of food-intake compensated by compulsive overeating (84, 90, 91), and amplify the saliency of food while the inhibitory control weakens (91). Decreased dopamine function is supported by studies showing declined D_2_R in obesity both in humans (92, 30, 84, 93) and in animals (90). However, in some studies these finding have not replicated, as the association between BMI and D_2_R was observed either positive and dependent on age (27) or nonexistent (94, 31, 83). Some studies also point towards a curvilinear relationship between BMI and D_2_R, such that the association is positive up to a certain BMI level after which the relation turns negative (95). The contribution of D_2_R genotype to obesity neither replicated in a large sample (96). The present large-scale study shows that the age-adjusted association of BMI and D_2_R availability is positive and linear, at least up to BMI of 30. It is thus possible that the effect is reversed beyond that point, but the current dataset does not have sufficient data for higher BMIs thus precluding such modeling that would be of great interest to confirm our finding. Overall, even though the estimates have some degree of uncertainty, we found no evidence for a negative relationship between BMI and striatal D_2_R availability.

### Lateralization of D_2_Rs

Lateralization effects, strongest in nucleus accumbens (right > left), were subtle with stronger hemispheric asymmetry of D_2_R availability in males than in females. Our finding of greater hemispheric asymmetry of males versus females may have resulted from better statistical power in the male sample and overall, the lateralization of D_2_Rs remains uncertain (97). However, some studies have found stronger left hemispheric lateralization of striatal D_2_R on preadolescent (98) and adult rats (99). In humans, meta-analyses suggest that emotion-related brain activity is more lateralized in males than females (62). Although the direction of lateralization is region-specific, it is consistently and similarly as in our data greater for men (62). Accordingly, sex differences in lateralized emotional processing in the brain may link with D_2_R expression, but this issue needs to be addressed in future studies. Finally, hemispheric asymmetry might relate to reward experiences involving dopaminergic function. Self-administered cocaine exposure evokes D_2_R lateralization (left > right) in male monkeys (100). In humans, evocative stimuli also elicit left lateralized brain activation, and in cocaine users (14 males, 3 females) it is particularly the cocaine-related cues that precede such activation (101). As cerebral lateralization and addictive behavior may both be more common in males, the interplay between these factors should be investigated in more detail.

### Limitations

The data were acquired using five different scanners. Although we adjusted for the differences between the scanners using statistical modeling, this may have introduced noise in the data. The predictor variables were not optimally balanced, with relatively high sex-ratio (120 males and 36 females) and different age ranges across sexes. We also did not have complete documentation about the reconstruction algorithms that have been used for all the studies, thus these could not be taken into account. While the reconstruction algorithms are typically stable for a particular scanner in our site, it is possible that several different reconstruction algorithms have been used for some of the scanners. Our statistical model was however flexible with respect to such variation, as the residual variances could vary by scanner.

In addition to age, sex and BMI, previous studies have revealed that genetic polymorphisms, such as A1 allele (79) and C957T (102), as well as other genetic and environmental factors that were not considered here may explain some of the individual differences in D_2_R availability (103). We used [11C]raclopride and BP_ND_ to measure striatal D_2_R availability. BP_ND_ being a product of receptor density and affinity (104) of unoccupied receptors (105), the level of binding reflects i) the D_2_R density, ii) the D_2_R affinity and ii) the D_2_R occupancy by endogenous dopamine (43, 45, 23, 106). Hence, using only one baseline image per subject we could not analyze receptor density and affinity separately (107). However, as the binding affinity between dopamine and D_2_R is assumed constant (108), as the endogenous dopamine does not override D_2_R antagonists (e.g. [11C]raclopride) as effectively as agonists (105) and as we used baseline scans including no interventions boosting dopamine firing (109), we expect the BP_ND_ to dominantly measure the D_2_R density.

### Conclusions

Striatal D_2_R availability decreases globally through age for both sexes. Females show on average 5-10% higher D_2_R availability than males. High BMI was associated with increased D_2_R availability, although this effect was only weak. D_2_R availability was more lateralized in males than in females, but the lateralization effects were overall subtle. Importantly, we confirmed that the template-based normalization method allows for accurate global ROI-level modeling of the PET data when deformation-field-based spatial normalization method is not possible due to missing MR image. In sum, D_2_R density is dependent on subject demographics, particularly on age and sex. These effects may contribute to age and sex dependent prevalence in neurological and psychiatric conditions involving altered D_2_R expression.

## Acknowledgements

This study was supported by the Academy of Finland (grants #294897 and #332225 to L.N.), the Sigrid Jusélius Foundation (grant to L.N.) and the Päivikki and Sakari Sohlberg Foundation (personal grant to T.M.). We thank senior researchers Marco Bucci and Semi Helin, as well as Professor Kari Auranen for sharing their expertise on kinetic modeling, radiochemistry and Bayesian data analysis, respectively.

