## Supplementary Material for "Age and sex dependent variability of type 2 dopamine receptors in the human brain: A large-scale PET cohort"

#### Scanner Considerations

The primary sample was imaged with five different scanners. The $BP_{ND}$ estimates clearly vary between the scanners (Figure S1), which was taken into account in the statistical modeling by adjusting for the scanner and allowing the scanner effect to vary regionally. The secondary sample was scanned with the five scanners presented here, as well as additional scanner ECAT.

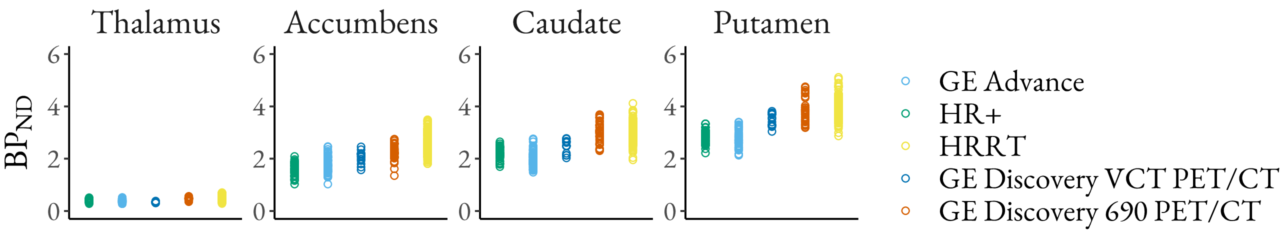


**Figure S1.** The $BP_{ND}$ estimates (original scale) for each scanner in each ROI in the primary sample. The figure shows that the $BP_{ND}$ estimates vary between the scanners and that the variation has regional differences.

While differences in how much the scanned individuals move during the data acquisition period could contribute to individual differences in $BP_{ND}$, we found no evidence for that (average framewise displacement used as a proxy for severity of movement).

#### Linearity Assessment of the Age and BMI effects

A recent article focusing on μ opioid receptors showed that the opioid receptor availability exhibits nonlinear associations with age in some brain regions (1). Thus, for the present sample we also estimated a nonlinear effect of age. Similarly, nonlinear effect was estimated for another continuous variable BMI (Figure S2), facilitating the detection of possible curvilinear association between BMI and $D_2R$ availability. The nonlinear effects were calculated separately for each ROI (single-ROI models). Hence, the only grouping variable used here was the scanner (compare scanner, ROI and their combination in the primary analysis). As in the primary modeling, the effects were calculated on a logarithmic scale.

*Age*

Assessing the linearity of age effect, the three out of four single-ROI models (putamen, caudate nucleus, nucleus accumbens) produced a few divergent transitions, suggesting poor fitness of the model. Hence, we adjusted the modeling settings to improve model fitness. As priors, we set both population-level effects (class = 'b') and SD (class = 'sd') as normally distributed with mean 0 and SD 0.5 (N (0,0.5)) (compare N (0,1) in the primary analysis). We set the AD as 0.999 (compare 0.99 in the primary analysis) (2, 3). These modifications improved the model fitness, leaving no divergent transitions. The Rhats were 1.

The results show that the age effect is well approximated by a linear function (Figure S2). As the number of subjects decreases toward the end of the age range, there is considerable uncertainty in the estimates of the ages above 60. To conclude, these analyses suggest that the association between age and [11C]raclopride $BP_{ND}$ is well approximated by a linear function.

*BMI*

We also estimated a nonlinear BMI effect (Figure S2). The model converged successfully (no divergent transitions, Rhats of 1) with primary settings: population-level effects (class = 'b') and SD (class = 'sd') as (N (0,1)) and AD as 0.99.

Similarly as with age, the BMI effect appears well approximated by a linear function, although the uncertainty increases toward the end of the BMI range, as a result of decreasing number of data points.

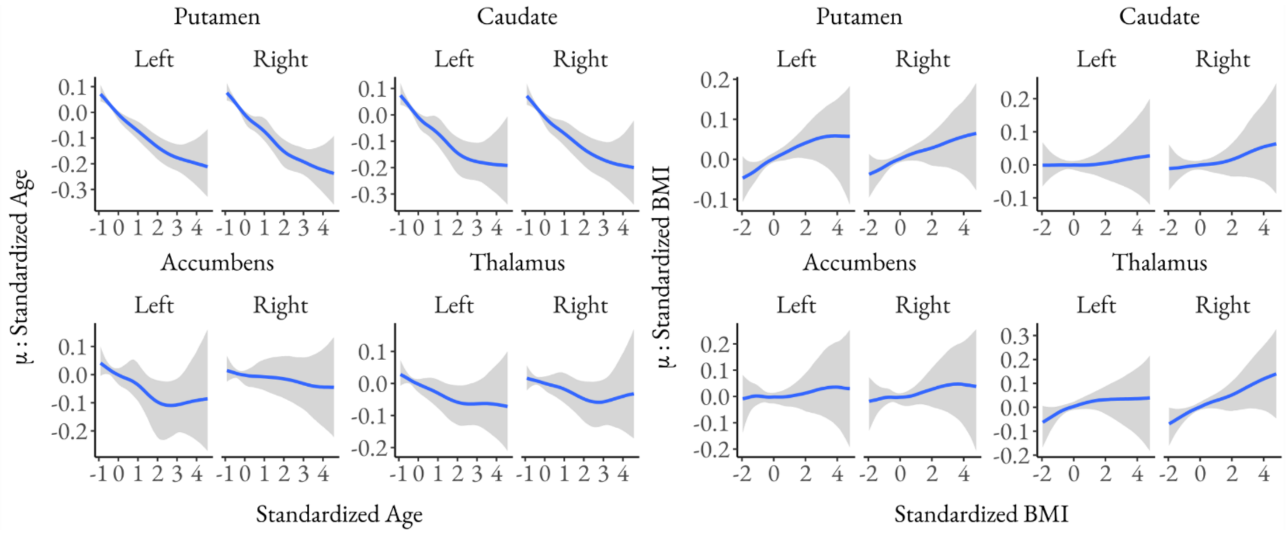


**Figure S2**. Assessment of the linearity of the age and BMI effects on $BP_{ND}$ (logarithmic scale) in each ROI, separately for the two hemispheres. The figure shows standardized age and BMI (x-axis) in relation to μ: standardized age and BMI (y-axis).

### Sampling Settings and Convergence Estimates

In the primary analysis, we used normal distributions with mean 0 and standard deviation (SD) 1 as priors for all (non-intercept) regression coefficients and half-normal distributions with mean 0 and SD 1 as priors for all SD parameters of the varying effects. On all other parameters, default priors of brms were used, which are defined to be non- or weakly informative (4). Markov chain Monte Carlo sampling was run with 5 chains, each with 4000 iterations (including 1000 warmups), adapt-delta (AD) of 0.99 and maximum treedepth of 20 (5, 6). The sampling produced no divergent transitions nor Rhats deviating from 1, supporting successful convergence of the chains (7). If not otherwise stated, these priors, sampling settings and convergence estimates (divergent transitions 0 and Rhats 1) are applicable to all modeling presented.

### Sensitivity Analysis

#### *Sensitivity analysis focusing on sex differences*

To assess if between-sex differences in the age distributions could explain the effects observed in the primary analysis, we executed a sensitivity analysis where we restricted the analysis to subjects less than 41 years of age, thus significantly balancing the age distributions between the sexes. See **Tables S1** and **S3**, as well as **Figure S3** for the sample characteristics. In a separate model, we additionally calculated the interaction of age and sex to validate that the age effect in the primary analysis could be calculated to the whole sample without separating males and females.

| | Males (n= 118) | | | Females (n= 22) | | |
|---|---|---|---|---|---|---|
| | Mean | SD | Range | Mean | SD | Range |
| Injected Activity (MBq) | 327 | 118 | 183-538 | 260 | 47 | 187-329 |
| Age (years) | 24 | 4 | 19-38 | 27 | 5 | 20-40 |
| Height (cm) | 181 | 7 | 167-199 | 166 | 5 | 156-174 |
| Weight (kg) | 79 | 12 | 58-130 | 58 | 5 | 48-66 |
| BMI (kg/m2) | 24 | 3 | 19-38 | 21 | 2 | 18-24 |

**Table S1.** Characteristics of the subsample (n= 140) aged 18-40.

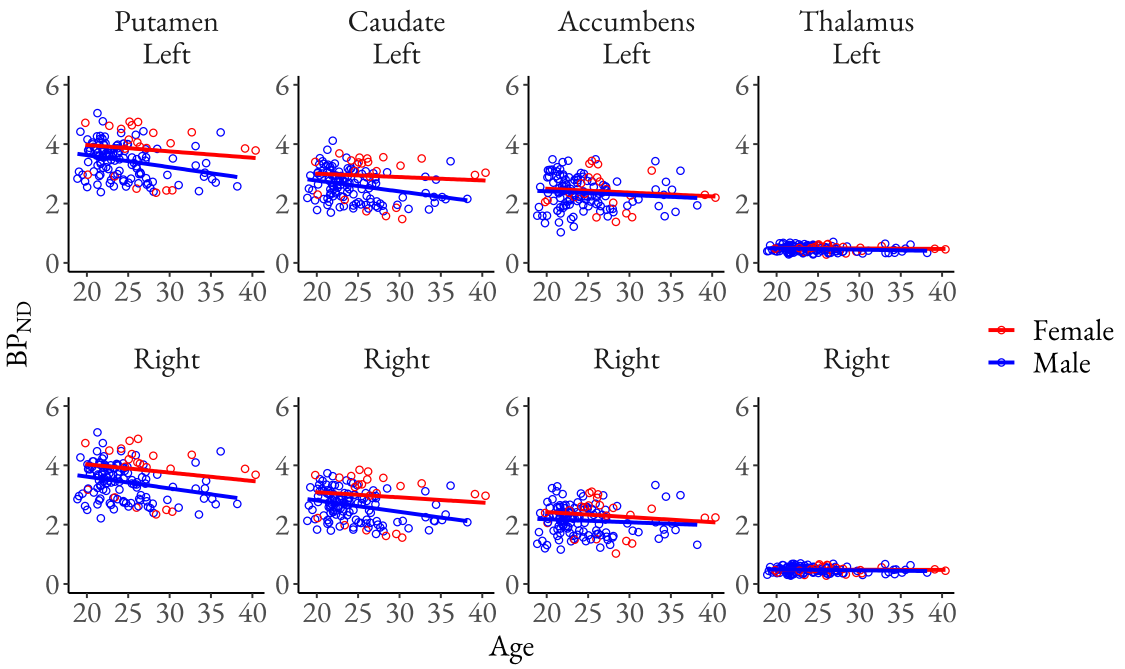


**Figure S3.** Left and right $D_2R$ $BP_{ND}$ (original scale) through age (original scale) in scatter plots separately for males and females in the primary dataset restricted to ages 40 and below. The figure shows the original $BP_{ND}$ estimates (points), linear regression lines separately for males and females (lines) and their 95% confidence intervals (shaded areas).

In the sample with age below 41 years, the results were essentially unchanged from the primary model. Hence, the sensitivity analysis validated the age, sex, BMI, and lateralization effects observed in the primary model (modeling results in Figures S4 and S5). On a separate analysis, the data did not show clear evidence for interaction of age and sex on $D_2R$ $BP_{ND}$ (modeling results in Figure S6). According to the data, the interactive relationship of age and sex still cannot completely be excluded, as globally the majority of the posterior uncertainty intervals are located above zero. However, the wide intervals are reflecting great uncertainty in the effect. Positive interaction effects would suggest steeper age-related decline in $BP_{ND}$ among males than females, coherently with previous literature reporting steeper $D_2R$ $BP_{ND}$ decrease of males versus females (8).

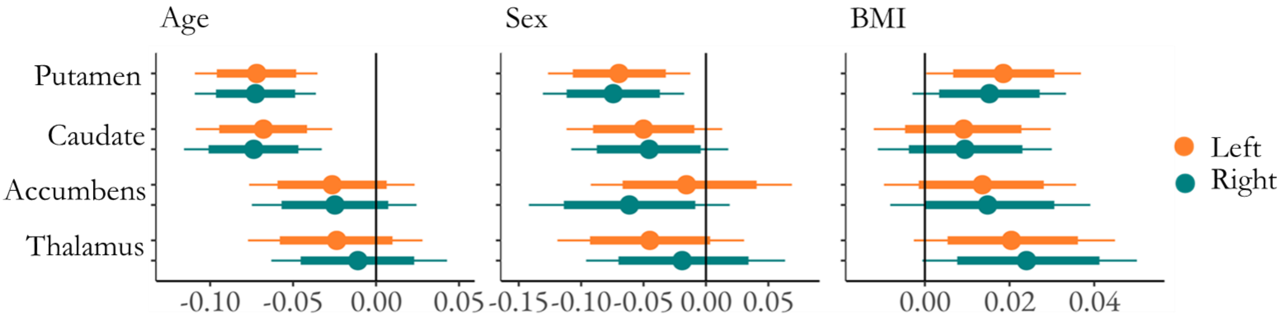


**Figure S4**. The effects of age (standardized), sex (male - female) and BMI (standardized) on $D_2R$ $BP_{ND}$ (logarithmic) in the primary model when the data was restricted to subjects aged 40 and below. The figure shows medians (circles), 80% (thick line) and 95% (thin line) posterior uncertainty intervals of the regression coefficients on a logarithmic scale.

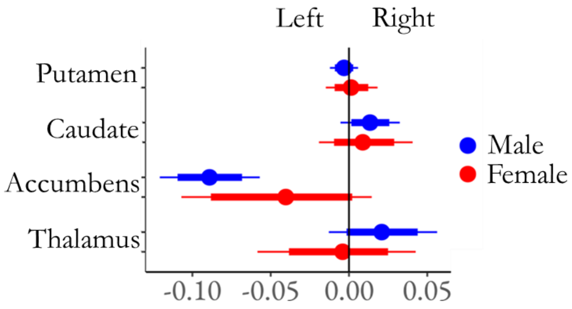


**Figure S5**. The effect of hemisphere (right - left) on $D_2R$ $BP_{ND}$ (logarithmic) in the primary model when the data was restricted to subjects aged 40 and below. The figure shows medians (circles), 80% (thick line) and 95% (thin line) posterior uncertainty intervals of the regression coefficients on a logarithmic scale.

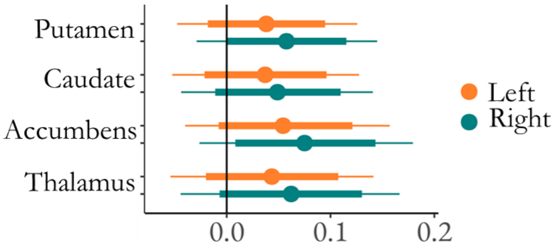


**Figure S6**. The interaction of age and sex (male - female) on $D_2R$ $BP_{ND}$ (logarithmic) in the subset restricted to subjects aged below 41 years. The figure shows medians (circles), 80% (thick line) and 95% (thin line) posterior uncertainty intervals of the regression coefficients on a logarithmic scale.

*Sensitivity analysis adjusting for regional volume*

We investigated whether adjusting for the volume of each ROI changes the results. The regional volumes were obtained by processing individual T1-weighted MR images with FreeSurfer (https://surfer.nmr.mgh.harvard.edu/). Prior to statistical modeling, the regional volume was standardized (i.e., z-scored) within each ROI to keep the regression coefficients on the same scale as all the other continuous predictors. To adjust for regional volume and to estimate its main effect on $BP_{ND}$, we added the volume as a new regressor in the primary model (along with age, sex, BMI and hemisphere).

Adjusting for the regional volumes did not change the overall results of age, sex, BMI or hemisphere (modeling results in Figures S7 and S8). The data neither showed clear and global evidence for the effects of regional volume itself, although in a single ROI, left accumbens, the posterior uncertainty interval did not overlap with zero (Figure S8), suggesting a positive association between the volume and $D_2R$ availability. The effect was positive, suggesting a 2% increase in $D_2R$ binding to co-occur with the increase of 1 SD (704 voxels, while mean 5209 voxels, voxel size dependent on the scanner, the mean across the scanners [1.3, 1.3, 2.9] mm) in the volume. Also, in bilateral caudate, the majority of the relatively narrow posterior uncertainty intervals located above zero, suggesting a positive effect also in caudate. Overall, we conclude that the main effects of regional volume may reflect the effect of partial volume correction.

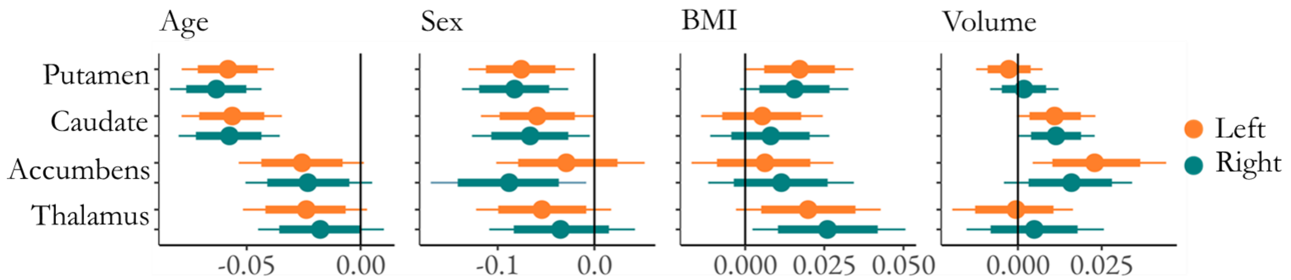


**Figure S7**. The effects of age (standardized), sex (male - female), BMI (standardized) and regional volume (standardized) on $D_2R$ $BP_{ND}$ (logarithmic) in the model that adjusted for regional volumes. The figure shows medians (circles), 80% (thick line) and 95% (thin line) posterior uncertainty intervals of the regression coefficients on a logarithmic scale.

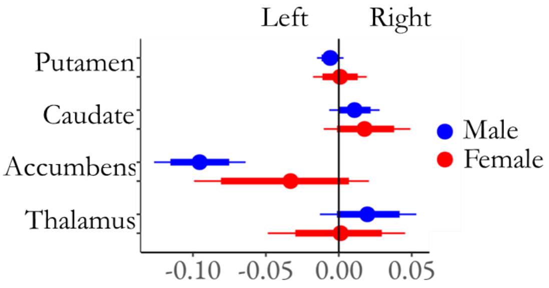


**Figure S8**. The effect of hemisphere (right - left) on $D_2R$ $BP_{ND}$ (logarithmic) in the model that adjusted for regional volumes. The figure shows medians (circles), 80% (thick line) and 95% (thin line) posterior uncertainty intervals of the regression coefficients on a logarithmic scale.

**Validation of an alternative approach for defining ROIs and reference regions**

We were able to identify altogether 304 baseline [11C]raclopride scans of healthy controls (including the primary sample of 156 scans). The deformation-field-based spatial normalization requiring high-resolution anatomical MR image was used as a primary normalization method for the primary sample. As for only 189 subjects the anatomical MR image was available, a total of 115 subjects had to be excluded from the primary sample due to missing MR image. Please see **Table S3** for the more detailed exclusion criteria.

To further validate our findings, we additionally used an alternative atlas and template-based normalization and modeling of the data to all identifiable 304 PET-scans, including those with missing MR image. The PET images were spatially normalized to an in-house [11C]raclopride template using SPM's "old normalize" tool. We used the Harvard-Oxford atlas to define regions of interest (nucleus accumbens, caudate nucleus, putamen and thalamus) as well as the cerebellar reference region.

We first validated the alternative atlas-based normalization approach by comparing the data produced by the two protocols. Hence, we used the sample that could be analyzed with both methods (i.e., subjects with the available MR image, n= 189, see Table S3). We found that the reference region time-activity curves generated by the alternative atlas-based approach corresponded closely with the time-activity curves of the primary FreeSurfer-based approach (Figure S9). Second, we found that the two methods produce highly correlative $BP_{ND}$ estimates in all ROIs (Figure S10). We also visually validated that the ROIs are well aligned. Overall, these results confirm that the template and atlas-based normalization yields reliable and accurate $BP_{ND}$ estimates.

In addition to validating that the alternative approach produces reliable and accurate $BP_{ND}$ estimates, we conducted additional statistical analysis to check whether our main findings - the effects of age and sex on $D_2R$ availability - would replicate in the secondary sample (see Table S3) that had been excluded from the primary dataset due to missing MR image or anthropometric information and normalized with the alternative method.

Similarly, as in the primary sample, the inclusion criteria for the secondary sample were that the injected dose was above 100 MBq and that only one scan (chronologically first) per subject was included in the analysis, if multiple baseline scans were available. In addition, from the secondary sample, five subjects had to be excluded because the field of view (FOV) was missing large parts of the cerebellum, making reliable reference-region-based quantification of tracer binding impossible. In addition, three observations from thalamus had to be excluded due to imprecise spatial normalization. These problematic observations were found during manually conducted visual quality control of the data. Characteristics of the secondary sample are presented in the **Table S2** and the exclusion criteria in the **Table S3**.

| | Males (n= 104) | | | Females (n= 31) | | |
|---|---|---|---|---|---|---|
| | Mean | SD | Range | Mean | SD | Range |
| Injected Activity (MBq) | 250 | 121 | 103-570 | 185 | 32 | 104-259 |
| Age (years) | 33 | 14 | 19-78 | 47 | 16 | 19-82 |

**Table S2.** Characteristics of the secondary sample (n= 135). SD= standard deviation.

In the secondary statistical analysis, we investigated the main effects of age and sex (but not BMI, as height and weight information was not available for all the subjects). We calculated the global ROI-level effects (i.e. not separately for left and right hemisphere) because the results from the primary analysis indicated that the age and sex effects are highly similar for the two hemispheres and because we only had available a [11C]raclopride template that had been averaged over the hemispheres, prohibiting reliable quantification of hemispheric differences. We first executed the secondary modeling with the same parameters and priors as in the primary analysis. However, the model produced 54 divergent transitions. Hence, we modified the parameters to improve the model fitness. We set the AD as 0.999. As priors, we set both population-level effects (class = 'b') and SD (class = 'sd') as normally distributed with mean 0 and SD 0.5 (N(0,0.5)) (2, 3). These modifications improved the model fitness. The divergent transitions decreased to 15 and all the Rhats were 1. The results shown in **Figure S11** are derived from the modified model with less divergent transitions.

The alternative normalization method replicated the main results from the primary analysis, supporting the decrease in $BP_{ND}$ through age and the increased $BP_{ND}$ of females versus males (Figure S11). Hence, this secondary analysis validated that the alternative normalization method can be used to estimate $BP_{ND}$ when the MR image is not available for the subject.

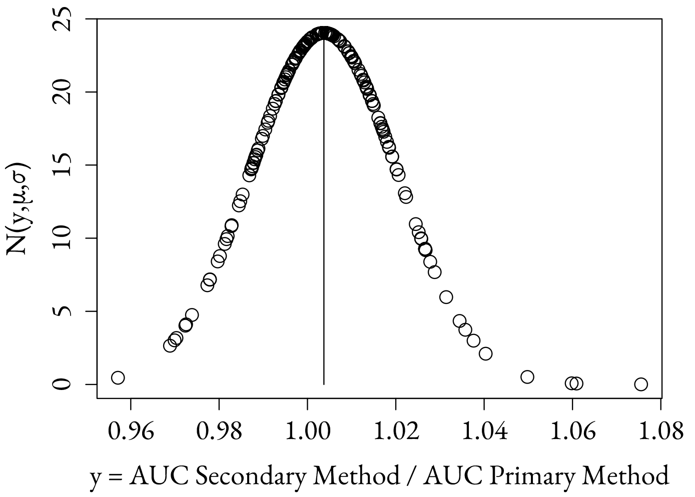


**Figure S9**. The figure demonstrates the ratios of the area under curve (AUC) estimates between the two spatial normalization methods in the reference area cerebellum for the subjects that MRI was available (n= 189). According to the figure, the two spatial normalization methods produce highly similar $BP_{ND}$ estimates (original scale) to the reference region, as most of the ratio observations are close to 1 (i.e., the AUCs are highly similar for both methods).

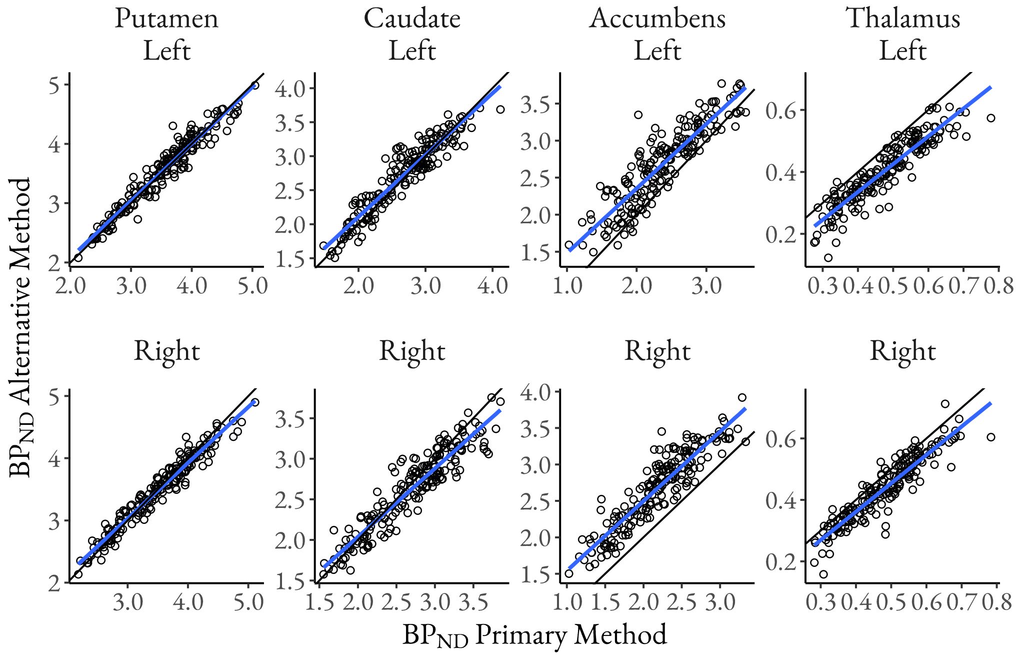


**Figure S10**. The scatter plots show the high correlation of the two different spatial normalization methods. On the x-axis, there are the $BP_{ND}$ estimates (original scale) acquired from the primary FreeSurfer based normalization method utilizing MR images. This was the normalization method used in the primary model. On the y-axis, there are the $BP_{ND}$ estimates (original scale) acquired with the alternative method, SPM's "old normalize" tool.

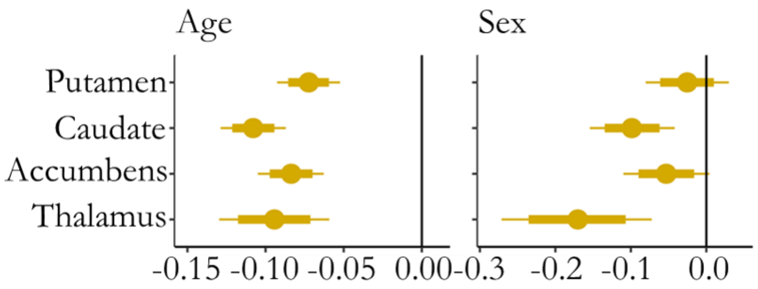


**Figure S11**. The global effects of age (standardized) and sex (male - female) on $D_2R$ $BP_{ND}$ (logarithmic) in the in the subset where the $BP_{ND}$ estimates were produced using the alternative spatial normalization method. The figure shows medians (circles), 80% (thick line) and 95% (thin line) posterior uncertainty intervals of the regression coefficients on a logarithmic scale.

| | Exclusion criteria | Primary sample | Subset aged 40 and under | Sample for validating normalization | Secondary sample |
|---|---|---|---|---|---|
| | MRI not available | 115 | 115 | 115 | NA |
| | Missing height or weight | 31 | 31 | NA | NA |
| | Dose missing or < 100 MBq | 2 | 2 | NA | 8 |
| | Suboptimal FOV | 0 | 0 | 0 | 5 |
| | Age above 40 | NA | 16 | NA | NA |
| | Inclusion in the primary sample | NA | NA | NA | 156 |
| Number of excluded from all (n= 304) | | 148 | 164 | 115 | 169 |
| Final sample size | | 156 | 140 | 189 | 135 |

**Table S3.** The table presents the exclusions made from each sample, starting from all identified [11C]raclopride chronologically first baseline scans of healthy subjects (n= 304). Sample for validating normalization= sample for calculating the correlation of the $BP_{ND}$ estimates produced by the two different normalization methods. Secondary sample= sample used for secondary data analysis testing the reproducibility of the age and sex effects. FOV= Field of view. The optimality of FOV was checked manually. NA= Not applicable, i.e. the exclusion criteria was not used for the sample.

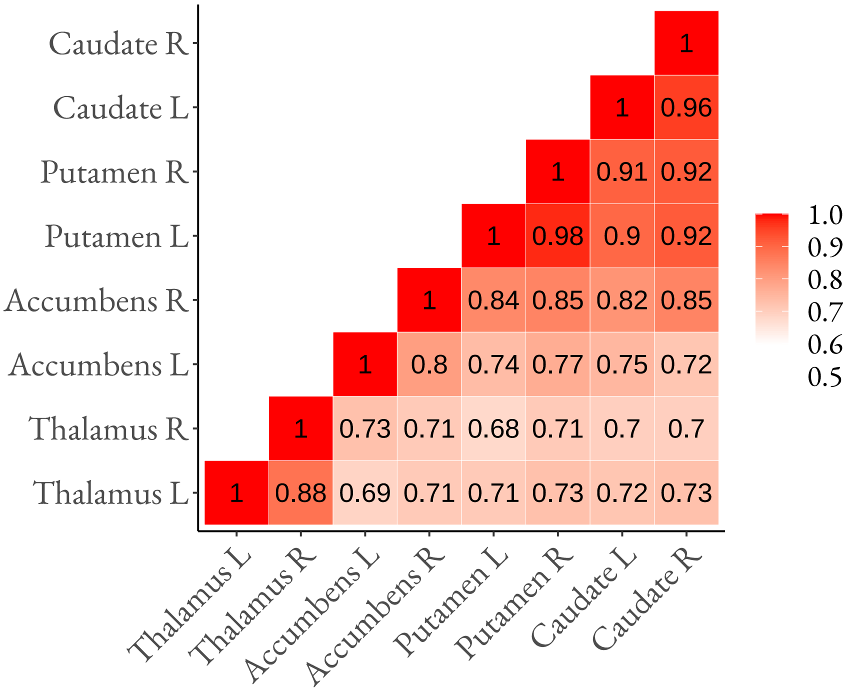


**Figure S12.** Between-ROI Pearson correlation coefficients of $BP_{ND}$ estimates (original scale) in the primary sample normalized with the primary method. The figure shows that the $BP_{ND}$ estimates correlate highly between the ROIs.
