## Supplementary Code for "Age and sex dependent variability of type 2 dopamine receptors in the human brain: A large-scale PET cohort"

5/17/2021

### Introduction

This is an R Markdown document describing the statistical modeling in our study investigating how age, sex, body mass index (BMI), as well as cerebral hemisphere and regional volume are associated with the type 2 dopamine receptor ($D_2R$) availability in the human brain.

### Subjects

The data were 156 baseline [11C]raclopride scans of healthy control subjects (120 males and 36 females; age 18-70 years, BMI range 17.7-38.0) scanned at Turku PET Centre between 2004 and 2018. Studies were included in the analysis if they were baseline scans with injected dose > 100 MBq (to avoid low signal-noise ratio (SNR)) and the magnetic resonance (MR) scan and basic demographic and anthropometric information (sex, age, height, weight) was available. If multiple baseline scans were acquired for a given individual, chronologically first scan was included in the analysis.

### Aims and Methods

In this cross-sectional register-based study we investigated how age, sex, BMI, cerebral hemisphere and regional volume influence $D_2R$ availability in the human brain using a large historical dataset (n=156) of [11C]raclopride PET scans performed between 2004 and 2018.

As regions of interest (ROI) we used striatal left (L) and right (R) hemisphere of putamen (put), caudate nucleus (cau), nucleus accumbens (acc) and extrastriatal thalamus (tha).

$D_2R$ availability was estimated from tracer binding in in $D_2Rs$. Tracer binding was quantified using the outcome measure binding potential ($BP_{ND}$), which is the ratio of specific binding to non-displaceable binding in tissue (Innis et al., 2007). ($BP_{ND}$) was estimated using a simplified reference tissue model (Lammertsma & Hume, 1996) with cerebellar gray matter serving as the reference region (Hall et al., 1996).

Statistical modeling was carried out in R (R Core Team, 2021) using brms (Bürkner, 2017, 2018) that applies the Markov-Chain Monte Carlo sampling tools of RStan (Stan Developent Team, 2020). The analysis script is available in Supplementary Material.

#### Load relevant libraries

```
library(ggplot2)
library(ggthemes)
library(bayesplot)
library(brms)
library(rstan)
library(ggpubr)
library(grid)
library(gridExtra)
library(ggcorrplot)
```

### Load required workspace

First, let's take a look at the $BP_{ND}$ estimates (original scale) for each ROI in the primary sample. The figure below shows medians (middle line), 25% (lower hinge) and 75% (upper hinge) quantiles, min value (lower whisker) and max value (upper whisker), as well as the data points for the original $D_2R$ $BP_{ND}$ estimates.

```
ggplot(data, aes(x=roi, y=bp))+
  geom_boxplot(fill = 'white')+
  geom_jitter(aes(colour = hemi),
              width= 0.1,
              pch=21)+
  scale_color_manual(labels= c("Left", "Right"),
                     values=c("#FF7F2A", "#008080"))+
  theme_classic()+
  labs(color = element_blank())+
  theme(axis.text.x=element_text(size=12),
        axis.text.y= element_text(size=12))+
  labs(y= expression(paste(BP[ND])),
       x= element_blank())+
  theme(legend.text= element_text(size=12))+
  theme(text=element_text(size=12))+
  scale_x_discrete(limits=c("tha", "nacc", "cau", "put"),
                   labels=(c("cau"= "Caudate",
                             "nacc"= "Accumbens",
                             "put"= "Putamen",
                             "tha"= "Thalamus")))
```

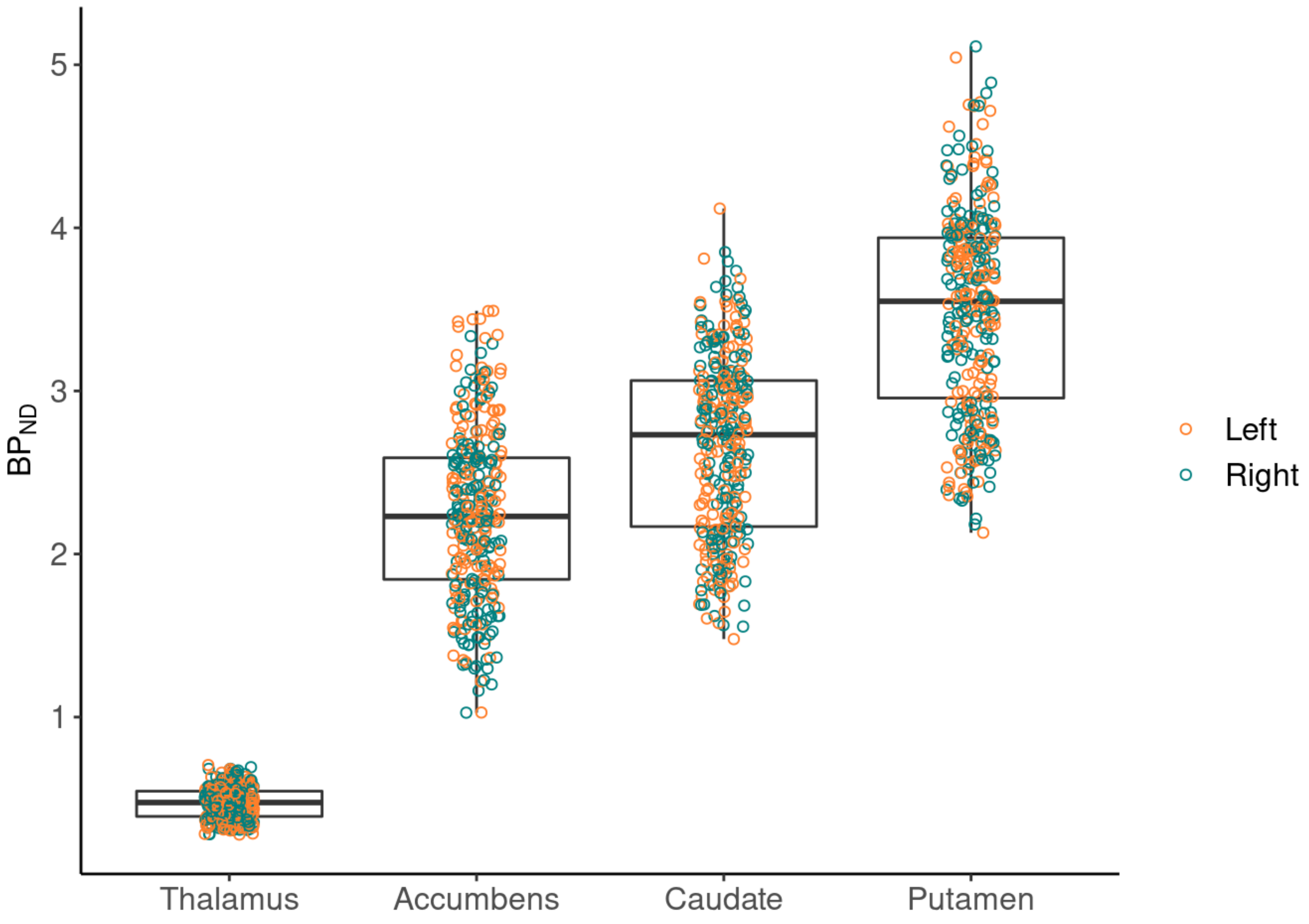


Next, let’s see the $BP_{ND}$ estimates (original scale) as a function of age (original scale). The figure shows the original $BP_{ND}$ estimates (points), linear regression lines separately for males and females (lines) and their 95% confidence intervals (shaded areas).

```
ggplot(data, aes(x=age, y=bp, color=sex))+
  geom_point(pch=21)+
  geom_smooth(method='lm')+
  ylim(0,6)+
  facet_wrap(~roihemi_f,
             labeller= labeller(roihemi_f = roihemi.labs),
             nrow = 2,
             scales = 'free') +
  theme_classic()+
  theme(strip.background = element_blank())+
  theme(legend.text= element_text(size=12))+
  theme(axis.text=element_text(size=12))+
  theme(strip.text=element_text(size=12))+
  theme(text=element_text(size=12))+
  labs(title= "", y=expression(paste(BP[ND])), x="Age", color = element_blank()) +
  scale_color_manual(labels= c("Female", "Male"), values= c("#FF0000", "#0000FF"))
+
  theme(aspect.ratio=1)+
  theme(plot.margin=unit(c(0,0,0,0), "cm"))
```

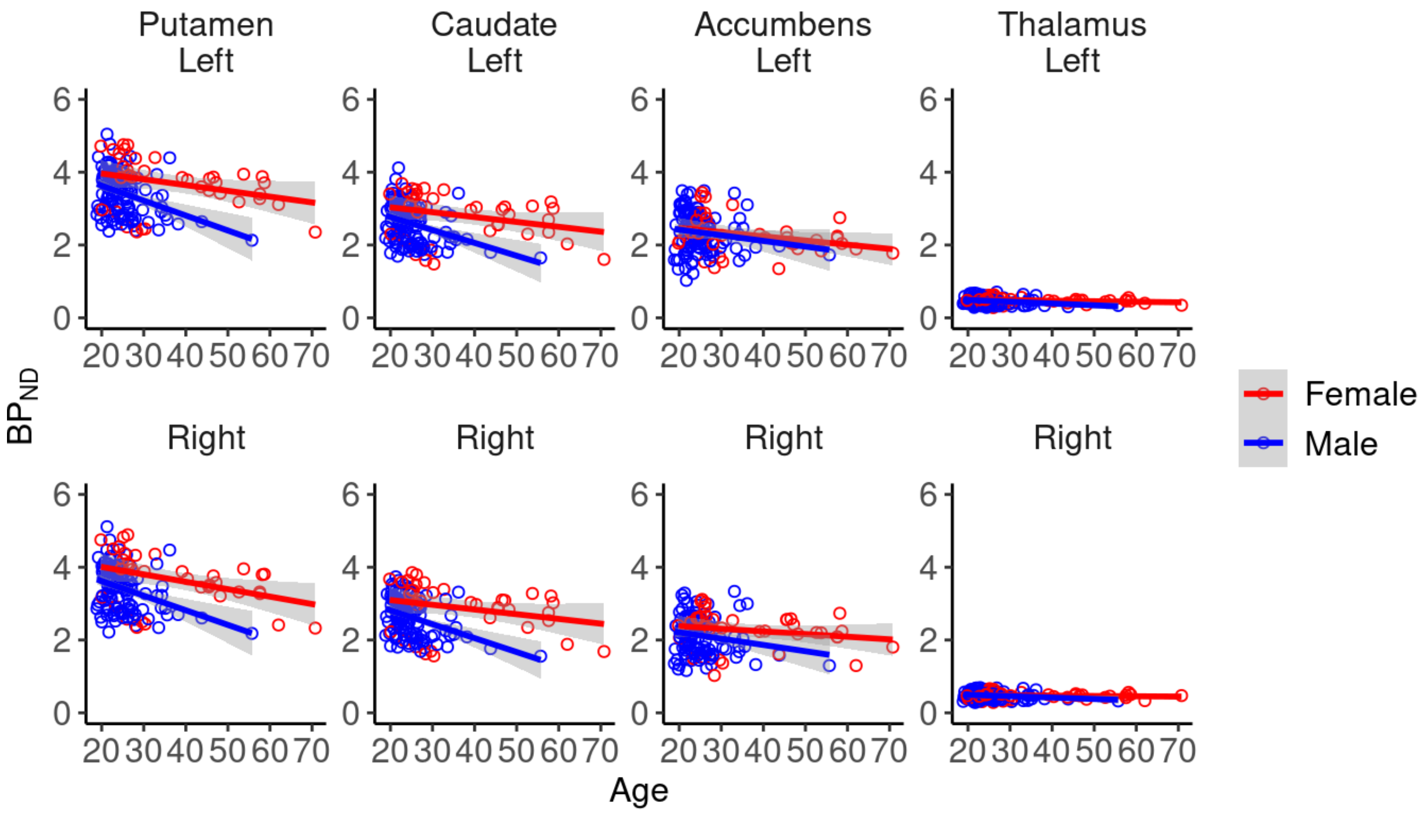


### Statistical Modeling: Primary Model

Next, the primary statistical modeling using brms is presented.

#### Model Settings: Primary Model

```
rstan_options(auto_write = TRUE)
ITER <- 4000
WARMUP <- 1000
NUM_CORES <- 5
CHAINS <- 5
AD <- 0.99
MAX_TREEDEPTH <- 20

custom_prior <- c(
  set_prior("normal(0,1)", class = "b"),
  set_prior("normal(0,1)", class = "sd")
)
```

#### Model: Primary Model

```
form.main <- bf(
  log_bp ~ (1 | subject) + (1 | gr(scanner, id = "scanner")) + (1 | gr(scanner:roi
, id = "scanner:roi")) + (1 + (age_z + sex + bmi_z)*hemi | gr(roi, id = "roi")) +
(age_z + sex + bmi_z)*hemi,
  sigma ~ (1 | gr(scanner, id = "scanner")) + (1 | gr(roi, id = "roi")) + (1 | gr(
scanner:roi, id = "scanner:roi"))
)

#fit.main <- brm(
#  formula = form.main,
#  data = data,
#  prior = custom_prior,
#  cores = NUM_CORES,
#  chains = CHAINS,
#  iter = ITER,
#  warmup = WARMUP,
#  control = list(adapt_delta = AD, max_treedepth = MAX_TREEDEPTH)
#)
```

### Results: Primary Model

Striatal $D_2R$ availability decreased through age for both sexes, and was higher in females (F) versus males (M) throughout the age range. BMI and striatal $D_2R$ availability were only weakly associated. There was no consistent lateralization of $D_2R$.

The below script shows an example of how to visualize the model results.

```
A1.main <- get_group_posteriors(fit.main, 'roi',c('age_z'))
colnames(A1.main)=c("nacc,age_z,left", "cau,age_z,left", "put,age_z,left", "tha,ag
e_z,left")
A2.main <- get_group_posteriors(fit.main, 'roi',c('age_z:hemir'))
A3.main <- A1.main + A2.main # create matrix
colnames(A3.main)= c("nacc,age_z,right", "cau,age_z,right",  "put,age_z,right", "t
ha,age_z,right")
A4.main <- cbind(A1.main, A3.main)

plotAmain <- mcmc_intervals(A4.main,
                            pars = c("tha,age_z,right",
                                     "tha,age_z,left",
                                     "nacc,age_z,right",
                                     "nacc,age_z,left",
                                     "cau,age_z,right",
                                     "cau,age_z,left",
                                     "put,age_z,right",
                                     "put,age_z,left"),
                            prob = 0.8, prob_outer = 0.95)+
  labs(title = "Age")+
  theme_classic()+
  scale_x_continuous(breaks= seq(-0.1,0.1, by= 0.05))+
  theme(text=element_text(size=12))+
  theme(axis.text=element_text(size=12))+
  geom_vline(xintercept = 0)+
  theme(plot.margin=unit(c(0,0,0,0), "cm"))+
  scale_y_discrete(labels=c("cau,age_z,right" = "Cau R",
                            "nacc,age_z,right"= "Acc R",
                            "put,age_z,right"="Put R",
                            "tha,age_z,right"="Tha R",
                            "cau,age_z,left" = "Cau L",
                            "nacc,age_z,left"= "Acc L",
                            "put,age_z,left"= "Put L",
                            "tha,age_z,left"= "Tha L"))
```

The figure below shows the effects of age (standardized), sex (male-female), BMI (standardized) on striatal and thalamic $D_2R$ binding estimates (logarithmic scale) separately for left and right hemisphere. In addition, the effect of hemisphere (right - left) is presented separately for males and females. The figure shows medians (circles), 80% (thick line) and 90% (thin line) posterior uncertainty intervals of the regression coefficients on a logarithmic scale.

```
ggarrange(plotAmain, plotBmain, plotCmain, plotDmain, ncol=2, nrow=2)
```

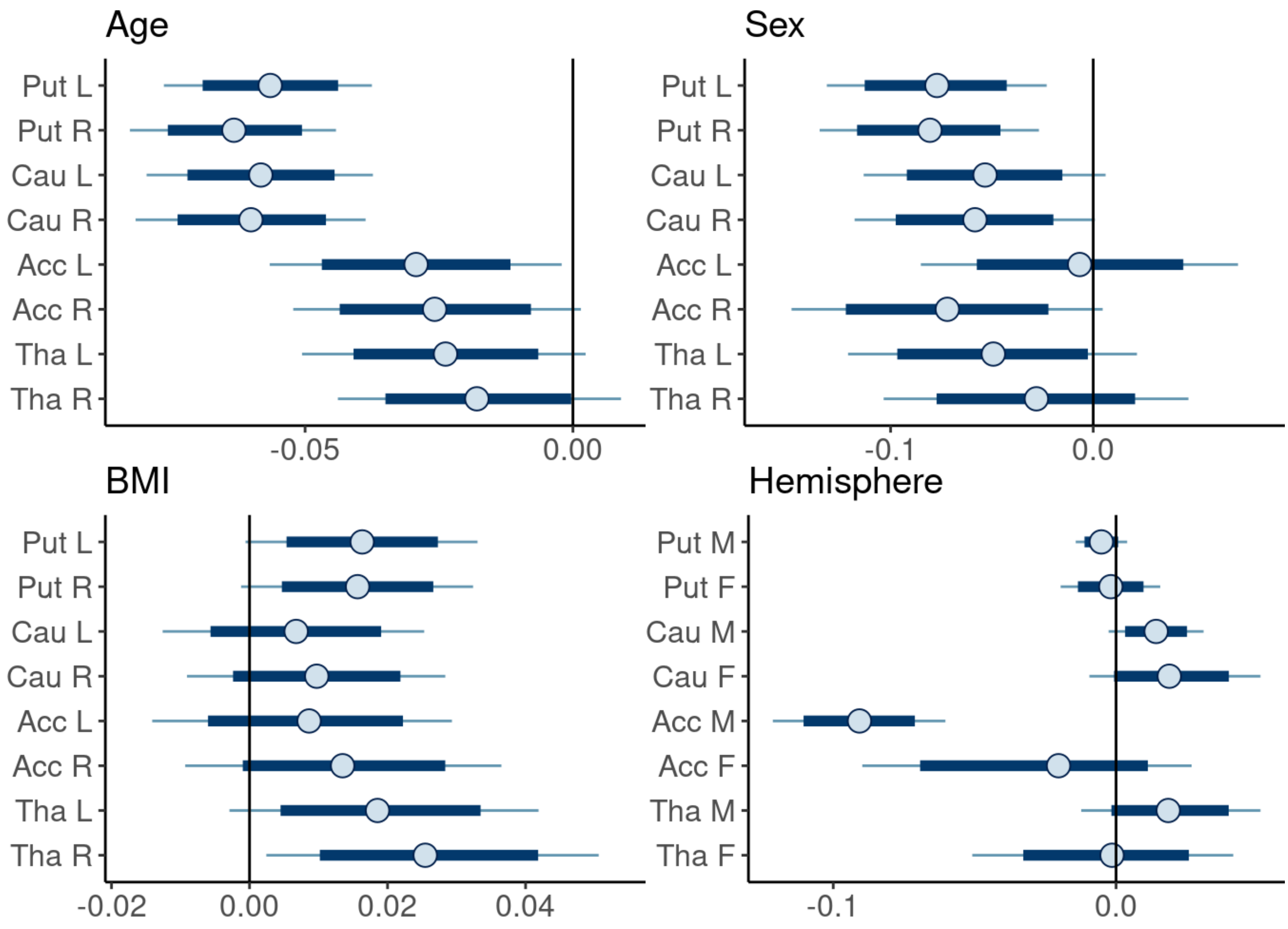


### Supplementary Material

#### Scanner Effect

Primary sample was scanned using five different scanners. The figure below presents the $BP_{ND}$ estimates (original scale) for each scanner in each ROI, showing that the $BP_{ND}$ estimates vary between the scanners and that the variation has regional differences.

```
ggplot(data, aes(x=reorder(scanner, bp, median), y=bp, color=scanner))+
  geom_point(pch=21)+
  ylim(0,6)+
  facet_wrap(~roi_f,
             labeller= labeller(roi_f = roi.labs),
             nrow = 2,
             scales = 'free') +
  theme_classic()+
  theme(strip.background = element_blank())+
  theme(text=element_text(size=14))+
  theme(axis.text.x = element_blank(), axis.ticks.x = element_blank())+
  labs(title= element_blank(), y=expression(paste(BP[ND])), x=element_blank(), col
or = element_blank()) +
  scale_color_manual(values= palette_pander(n=5),
                      labels = c("GE Advance", "HR+", "HRRT", "GE Discovery VCT PET
/CT", "GE Discovery 690 PET/CT"))+
  theme(legend.position = "right")+
  theme(legend.text=element_text(size=12),
        strip.text.x = element_text(size=14))+
  theme(aspect.ratio=1)+
  theme(plot.margin=unit(c(0,0,0,0), "cm"))
```

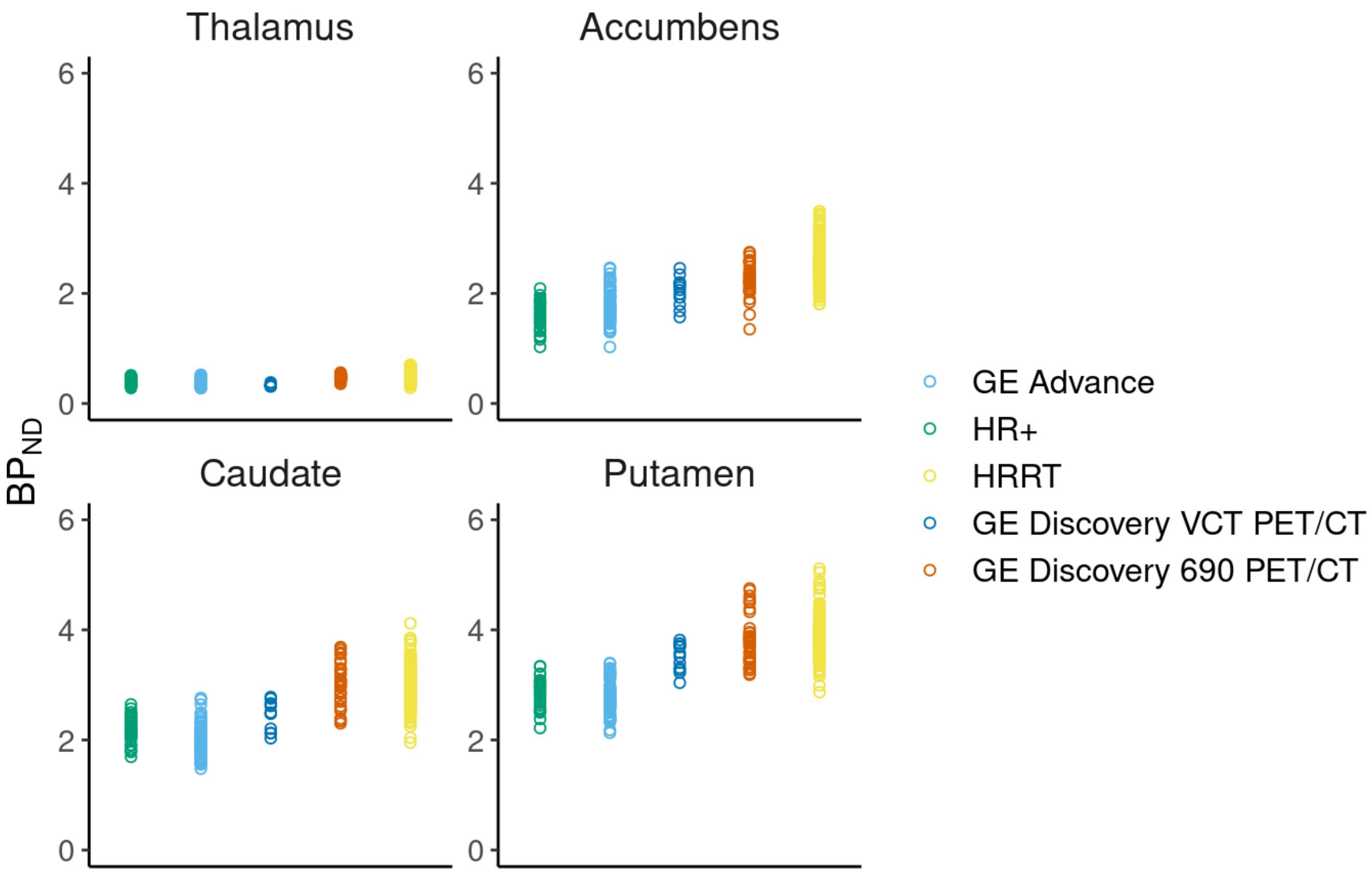


#### Linearity Assessment of Age and BMI Effects

The linearity assessment of the continous variables age and BMI was done by executing a model that allowed nonlinearity of the effects. The modeling was done separately for each ROI (single ROI models).

Using the same model settings as in the primary model, modeling the age effects produced a few divergent transitions. Hence, we modified the settings to improve model fitness (please see modeling settings below). The models assessing BMI effects, however, were executed with the same settings as in the primary model and such modeling produced no divergent transitions (see the script below). The script below demonstrates the modeling conducted for each ROI.

#### Model: Linearity of the Age and BMI Effects

###### Age

```
rstan_options(auto_write = TRUE)
ITER <- 4000
WARMUP <- 1000
NUM_CORES <- 5
CHAINS <- 5
AD <- 0.999 # modified from 0.99
MAX_TREEDEPTH <- 20

custom_prior <- c(
  set_prior("normal(0,0.5)", class = "b"), # modified from normal(0,1)
  set_prior("normal(0,0.5)", class = "sd") # modified from normal(0,1)
)

form.NLage.nacc.mod <- bf(
  log_bp ~ (1 | gr(scanner, id = "scanner")) + s(age_z, by = hemi) + sex + bmi_z,
  sigma ~ (1 | gr(scanner, id = "scanner"))
)

#fit.NLage.nacc <- brm(
#  formula = form.NLage.nacc,
#  data = data[data$roi=='nacc',],
#  prior = custom_prior,
#  cores = NUM_CORES,
#  chains = CHAINS,
#  iter = ITER,
#  warmup = WARMUP,
#  control = list(adapt_delta = AD, max_treedepth = MAX_TREEDEPTH)
#)
```

###### BMI

```
rstan_options(auto_write = TRUE)
ITER <- 4000
WARMUP <- 1000
NUM_CORES <- 5
CHAINS <- 5
AD <- 0.99
MAX_TREEDEPTH <- 20

custom_prior <- c(
  set_prior("normal(0,1)", class = "b"),
  set_prior("normal(0,1)", class = "sd")
)

form.NLbmi.nacc <- bf(
  log_bp ~ (1 | gr(scanner, id = "scanner")) + age_z + sex + s(bmi_z, by = hemi),
  sigma ~ (1 | gr(scanner, id = "scanner"))
)

#fit.NLbmi.nacc <- brm(
#  formula = form.NLbmi.nacc,
#  data = data[data$roi=='nacc',],
#  prior = custom_prior,
#  cores = NUM_CORES,
#  chains = CHAINS,
#  iter = ITER,
#  warmup = WARMUP,
#  control = list(adapt_delta = AD, max_treedepth = MAX_TREEDEPTH)
#)
```

##### Results: Linearity of the Age and BMI Effects

The results show that the age and BMI effects on a logarithmic scale are very well approximated by a linear function. As the number of subjects decreases toward the end of the age range, there is considerable uncertainty in the estimates of the ages above 60. To conclude, these analyses suggest that the association between age and BMI with log-transformed [11C]raclopride $BP_{ND}$ is well approximated by a linear function at least up to age 60.

The below script and figures show an example of how to visualize the modeling results. The figures below show linearity assessment of the age and BMI effects in each ROI, separately for the two hemispheres. On the x-axis is the standardized age and on the y-axis the mu:nonlinear age.

```
#nla1mod.Rmd <- plot(conditional_smooths(fit.NLage.put.mod))[[1]]+
#  theme_classic()+
#  theme(strip.background = element_blank())+
#  theme(text=element_text(size=12))+
#  labs(title="Putamen", x =element_blank(), y = element_blank())+
#  facet_grid(.~ hemi, labeller = labeller(hemi = hemi.labs))+
#  theme(strip.text=element_text(size=12))+
#  theme(axis.text=element_text(size=12))+
#  theme(plot.margin=unit(c(0,0,0,0), "cm"),
#        plot.title=element_text(size= 12, hjust=0.5))
```

```
grid.arrange(nla1mod.Rmd, nla2mod.Rmd, nla3mod.Rmd, nla4mod.Rmd, nrow= 2,
                    left = textGrob(label = expression(paste(mu," : Standardized A
ge")), rot = 90, gp=gpar(fontsize = 12, vjust = 1)),
                    bottom = textGrob(label ="Standardized Age", gp=gpar(fontsize
= 12, vjust = 1)),
                    vp= viewport(width=0.99, height=0.99))
```

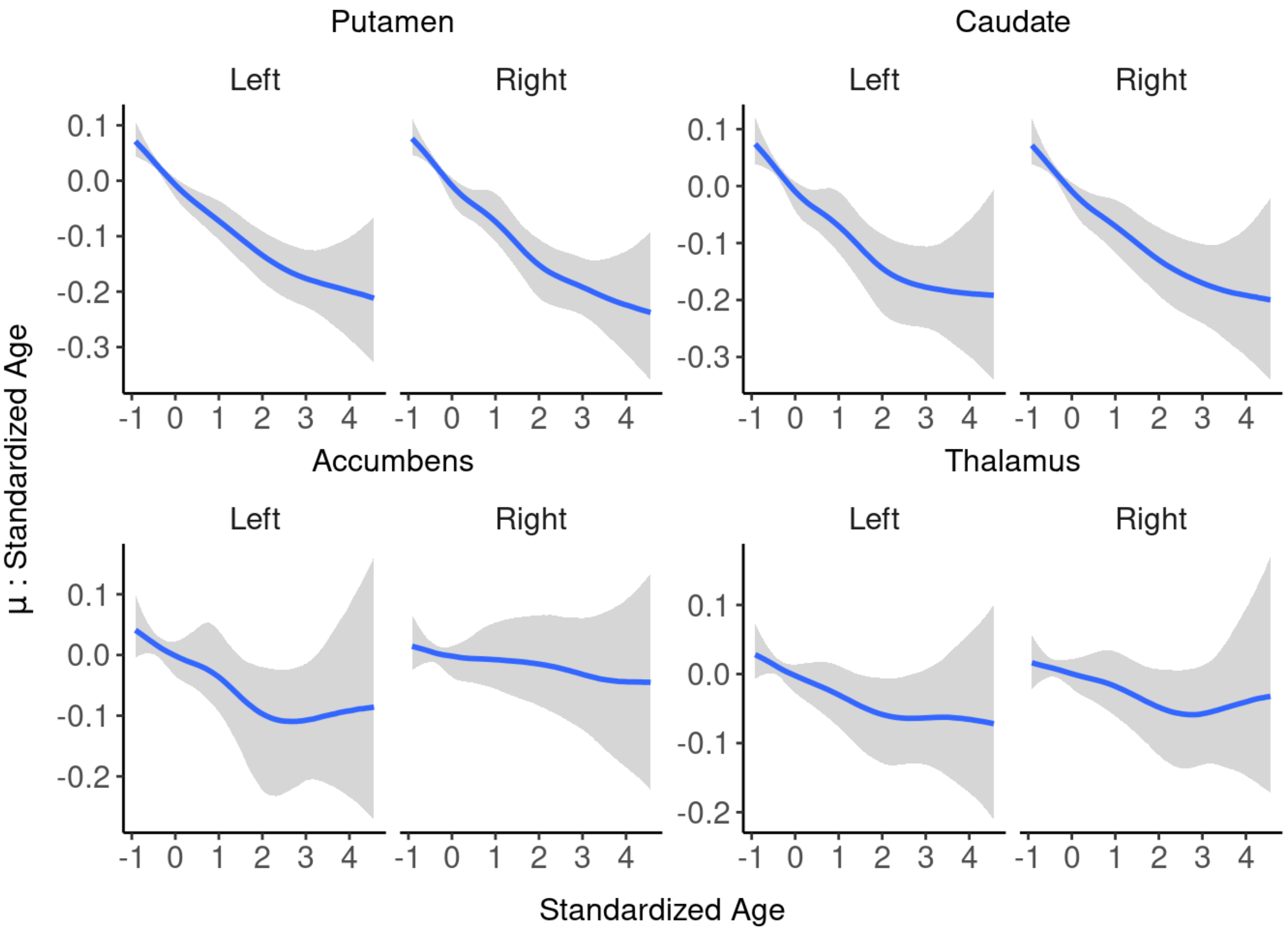


```
grid.arrange(nlb1.Rmd, nlb2.Rmd, nlb3.Rmd, nlb4.Rmd, nrow= 2,
                    left = textGrob(label = expression(paste(mu," : Standardized B
MI")), rot = 90, gp=gpar(fontsize = 12, vjust = 1)),
                    bottom = textGrob(label ="Standardized BMI", gp=gpar(fontsize
= 12, vjust = 1)),
                    vp= viewport(width=0.99, height=0.99))
```

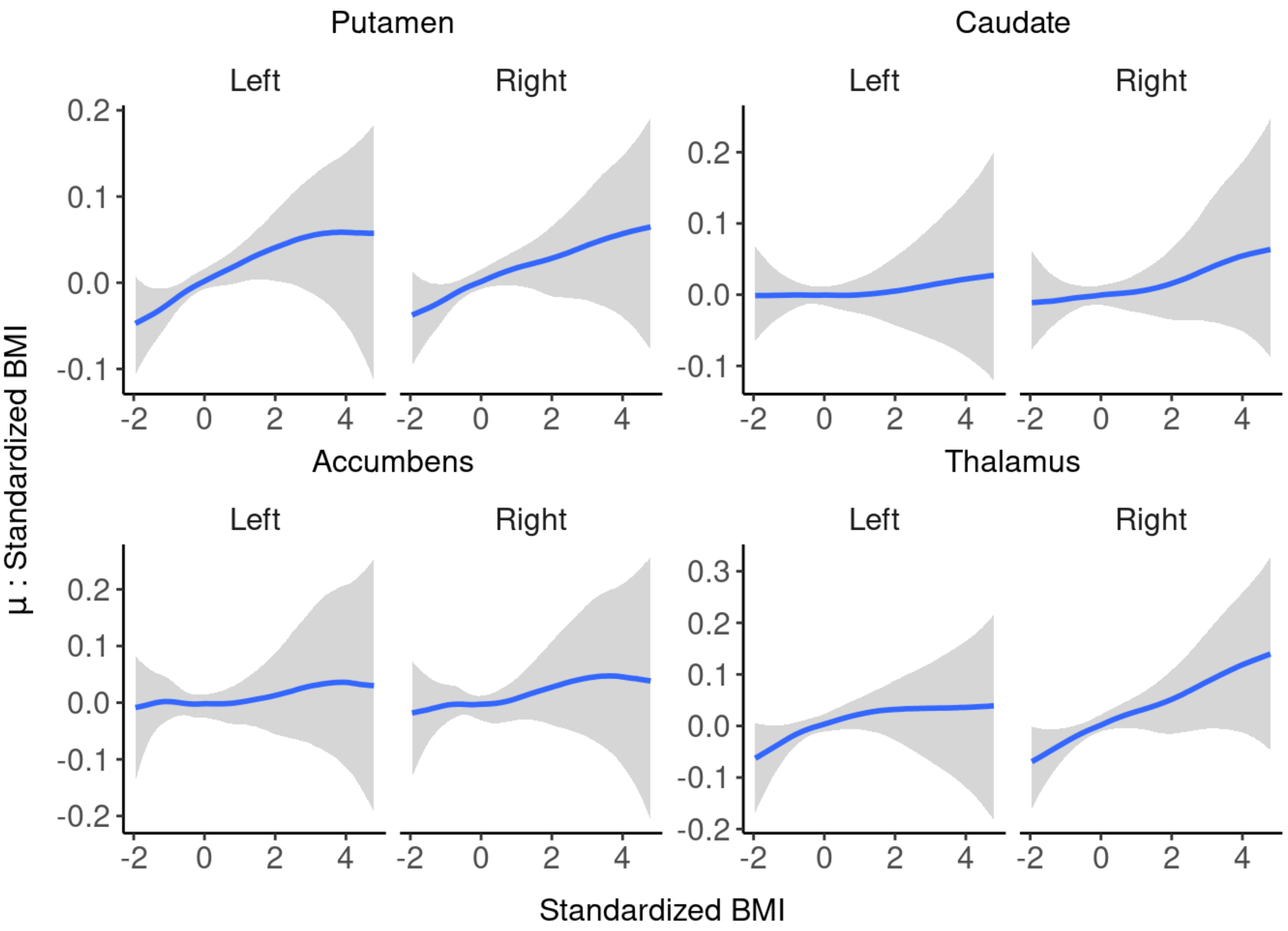


### Sensitivity Analysis

#### Sensitivity Analysis Focusing on Sex Differeces

To assess if between-sex differences in the age distributions could explain the observed effects, we executed a sensitivity analysis where we restricted the analysis to subjects less than 41 years of age, thus significantly balancing the age distributions between the sexes.

Let's see the $BP_{ND}$ estimates (original scale) as a function of age (original scale). The figure shows the original $BP_{ND}$ estimates (points), linear regression lines separately for males and females (lines) and their 95% confidence intervals (shaded areas).

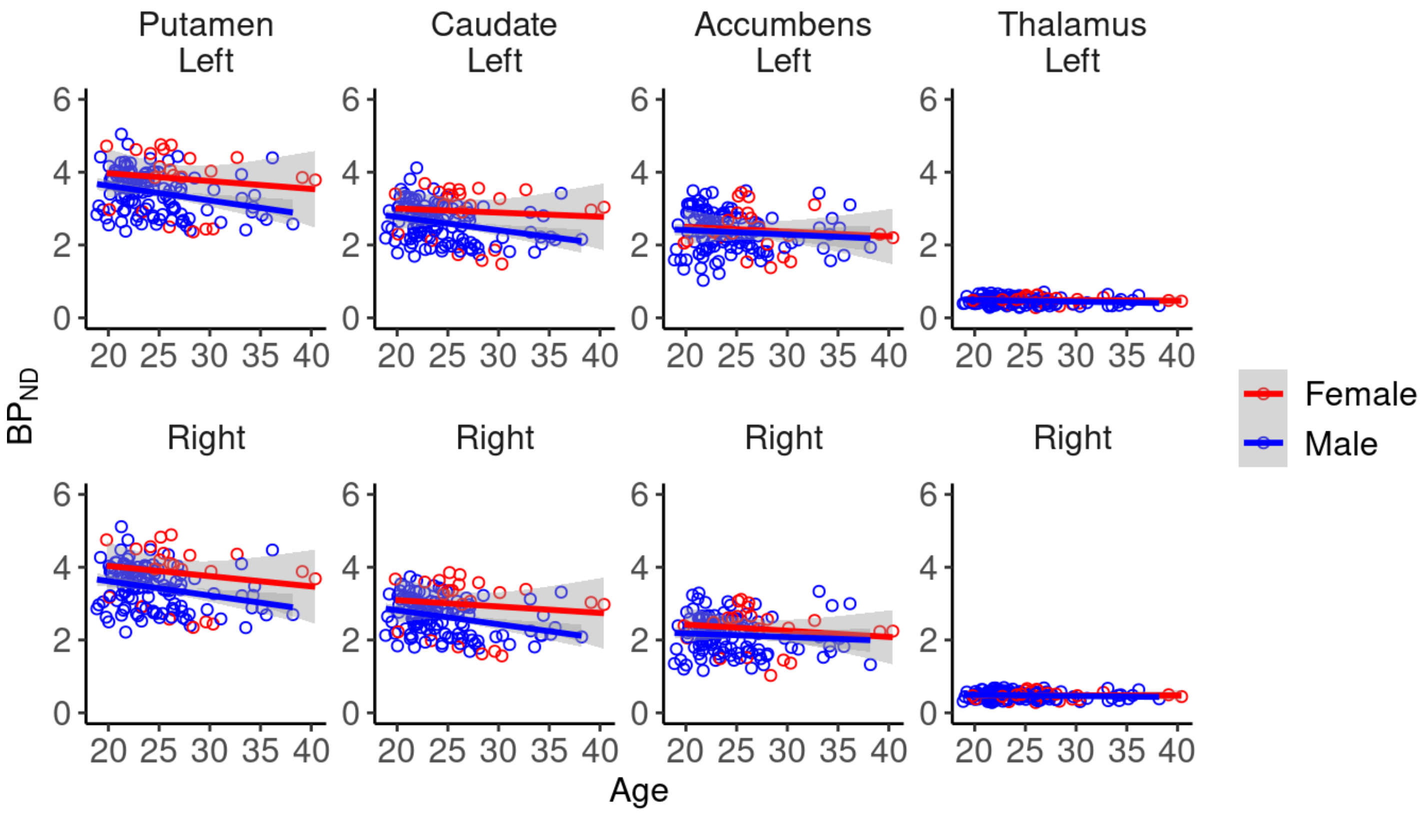


Let's then take a look at the model.

##### Model: Subjects Aged 18-40

```
form.40 <- bf(
  log_bp ~ (1 | subject) + (1 | gr(scanner, id = "scanner")) + (1 | gr(scanner:roi
, id = "scanner:roi")) + (1 + (age_z + sex + bmi_z)*hemi | gr(roi, id = "roi")) +
(age_z + sex + bmi_z)*hemi,
  sigma ~ (1 | gr(scanner, id = "scanner")) + (1 | gr(roi, id = "roi")) + (1 | gr(
scanner:roi, id = "scanner:roi"))
)

#fit.40 <- brm(
#  formula = form.40,
#  data = data_40,
#  prior = custom_prior,
#  cores = NUM_CORES,
#  chains = CHAINS,
#  iter = ITER,
#  warmup = WARMUP,
#  control = list(adapt_delta = AD, max_treedepth = MAX_TREEDEPTH)
#)
```

##### Results: Subjects Aged 18-40

The figure below shows, in the primary model when the data was restricted to subjects aged 40 and below, the effects of age (standardized), sex (male-female), BMI (standardized) on striatal and thalamic $D_2R$ binding (logarithmic scale) separately for left and right hemisphere. In addition, the effect of hemisphere (right - left) is presented separately for males and females. The figure shows medians (circles), 80% (thick line) and 90% (thin line) posterior uncertainty intervals of the regression coefficients on a logarithmic scale.

```
ggarrange(plotA40, plotB40, plotC40, plotD40, ncol= 2, nrow = 2)
```

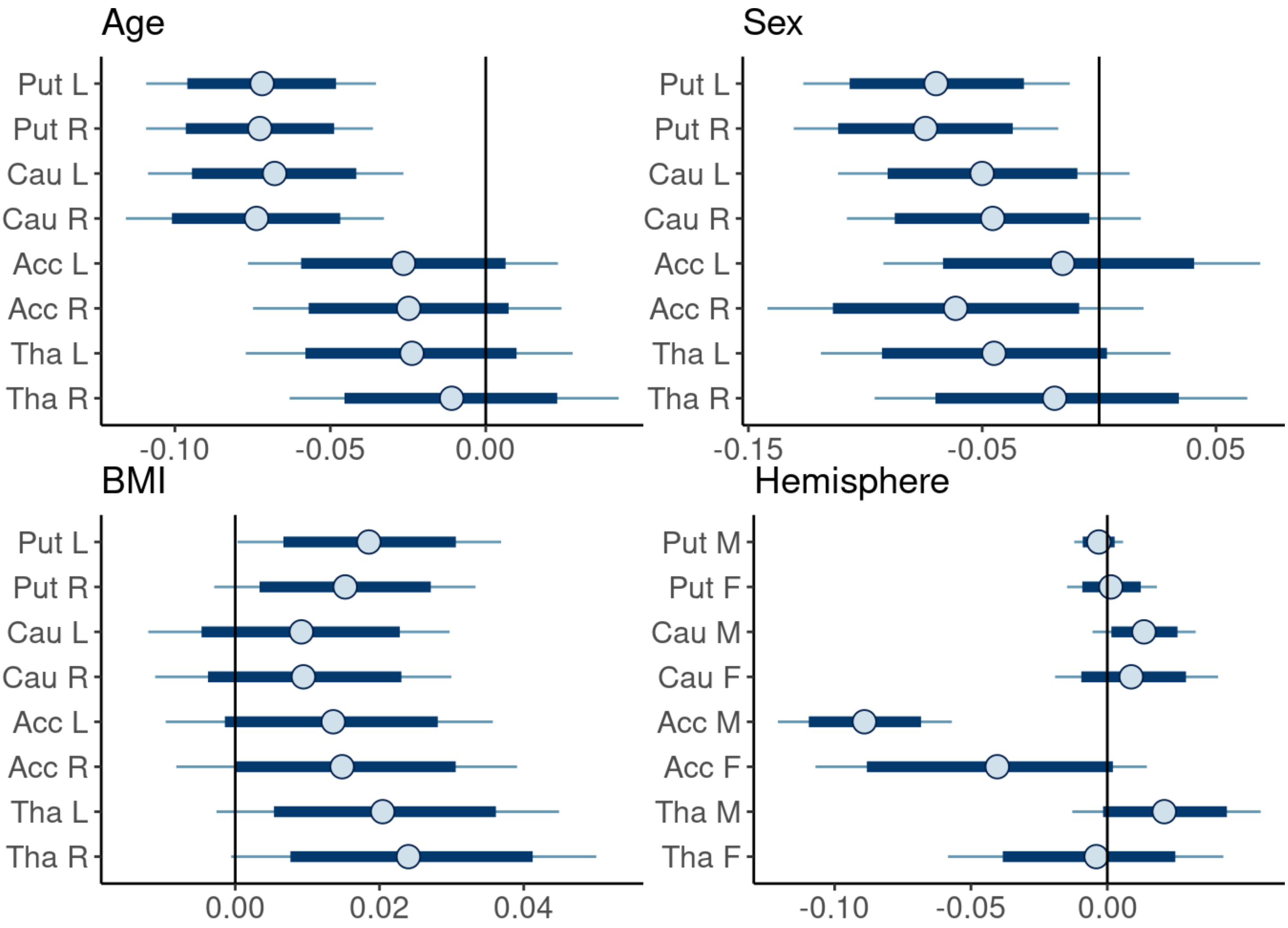


In a separate model, we additionally calculated the interaction of age and sex to validate that the age effect in the primary analysis could be calculated to the whole sample and not separately for males and females.

##### Model: Sex-Age Interaction in Subjects Aged 18-40

```
form.40.i <- bf(
  log_bp ~ (1 | subject) + (1 | gr(scanner, id = "scanner")) + (1 | gr(scanner:roi
, id = "scanner:roi")) + (1 + (age_z*sex + bmi_z)*hemi | gr(roi, id = "roi")) + (a
ge_z*sex + bmi_z)*hemi,
  sigma ~ (1 | gr(scanner, id = "scanner")) + (1 | gr(roi, id = "roi")) + (1 | gr(
scanner:roi, id = "scanner:roi"))
)

#fit.40.i <- brm(
#  formula = form.40.i,
#  data = data_40,
#  prior = custom_prior,
#  cores = NUM_CORES,
#  chains = CHAINS,
#  iter = ITER,
#  warmup = WARMUP,
#  control = list(adapt_delta = AD, max_treedepth = MAX_TREEDEPTH)
#)
```

##### Results: Sex-Age Interaction in Subjects Aged 18-40

The figure below shows the interaction of age and sex (male - female) on striatal and thalamic $D_2R$ binding (logarithmic scale) separately for left and right hemisphere in the subset restricted to subjects aged below 41 years. The figure shows medians (circles), 80% (thick line) and 90% (thin line) posterior uncertainty intervals of the regression coefficients on a logarithmic scale.

```
mcmc_intervals(AS4.40.i,
               pars = c("tha,age_z:sexm,right",
                        "tha,age_z:sexm,left",
                        "nacc,age_z:sexm,right",
                        "nacc,age_z:sexm,left",
                        "cau,age_z:sexm,right",
                        "cau,age_z:sexm,left",
                        "put,age_z:sexm,right",
                        "put,age_z:sexm,left"),
               prob = 0.8, prob_outer = 0.95)+
  labs(title = "Sex-Age Interaction")+
  theme_classic()+
  theme(text=element_text(size=12))+
  theme(axis.text=element_text(size=12))+
  geom_vline(xintercept = 0)+
  theme(plot.margin=unit(c(0,0,0,0), "cm"))+
  scale_x_continuous(breaks= seq(-0.1,0.5, by= 0.05))+
  scale_y_discrete(labels=(c("tha,age_z:sexm,right" = "Tha R",
                             "tha,age_z:sexm,left" = "Tha L",
                             "nacc,age_z:sexm,right" = "Acc R",
                             "nacc,age_z:sexm,left" = "Acc L",
                             "cau,age_z:sexm,right" = "Cau R",
                             "cau,age_z:sexm,left" = "Cau L",
                             "put,age_z:sexm,right" = "Put R",
                             "put,age_z:sexm,left" = "Put L")))
```

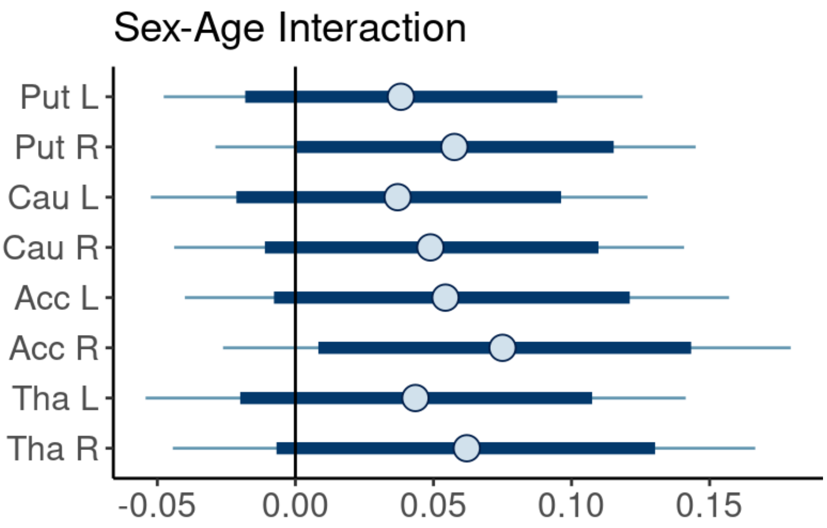


##### Sensitivity Analysis Adjusting for Regional Volume

To adjust for regional cerebral volume and to estimate its own main effect on $BP_{ND}$, we added the the volume as a new regressor in the primary model (along with age, sex, BMI and hemisphere). Adjusting for the regional volumes did not change the overall results of age, sex, BMI or hemisphere. However, the data suggested that the regional volume itself was connected to $BP_{ND}$ in left accumbens, where posterior uncertainty interval did not overlap with zero. However, the main effects of regional volume may reflect the effect of partial volume correction.

###### Model: Regional Volume

```
form.vol <- bf(
  log_bp ~ (1 | subject) + (1 | gr(scanner, id = "scanner")) + (1 | gr(scanner:roi
, id = "scanner:roi")) + (1 + (age_z + sex + bmi_z + vol_z)*hemi | gr(roi, id = "r
oi")) + (age_z + sex + bmi_z + vol_z)*hemi,
  sigma ~ (1 | gr(scanner, id = "scanner")) + (1 | gr(roi, id = "roi")) + (1 | gr(
scanner:roi, id = "scanner:roi"))
)

#fit.vol <- brm(
#  formula = form.vol,
#  data = data,
#  prior = custom_prior,
#  cores = NUM_CORES,
#  chains = CHAINS,
#  iter = ITER,
#  warmup = WARMUP,
#  control = list(adapt_delta = AD, max_treedepth = MAX_TREEDEPTH)
#)
```

###### Results: Model with Regional Volume

The figure below shows the effects of age (standardized), sex (male-female), BMI (standardized) on striatal and thalamic $D_2R$ binding (logarithmic scale) separately for left and right hemisphere in the model that adjusted for regional volumes. The figure shows medians (circles), 80% (thick line) and 90% (thin line) posterior uncertainty intervals of the regression coefficients on a logarithmic scale.

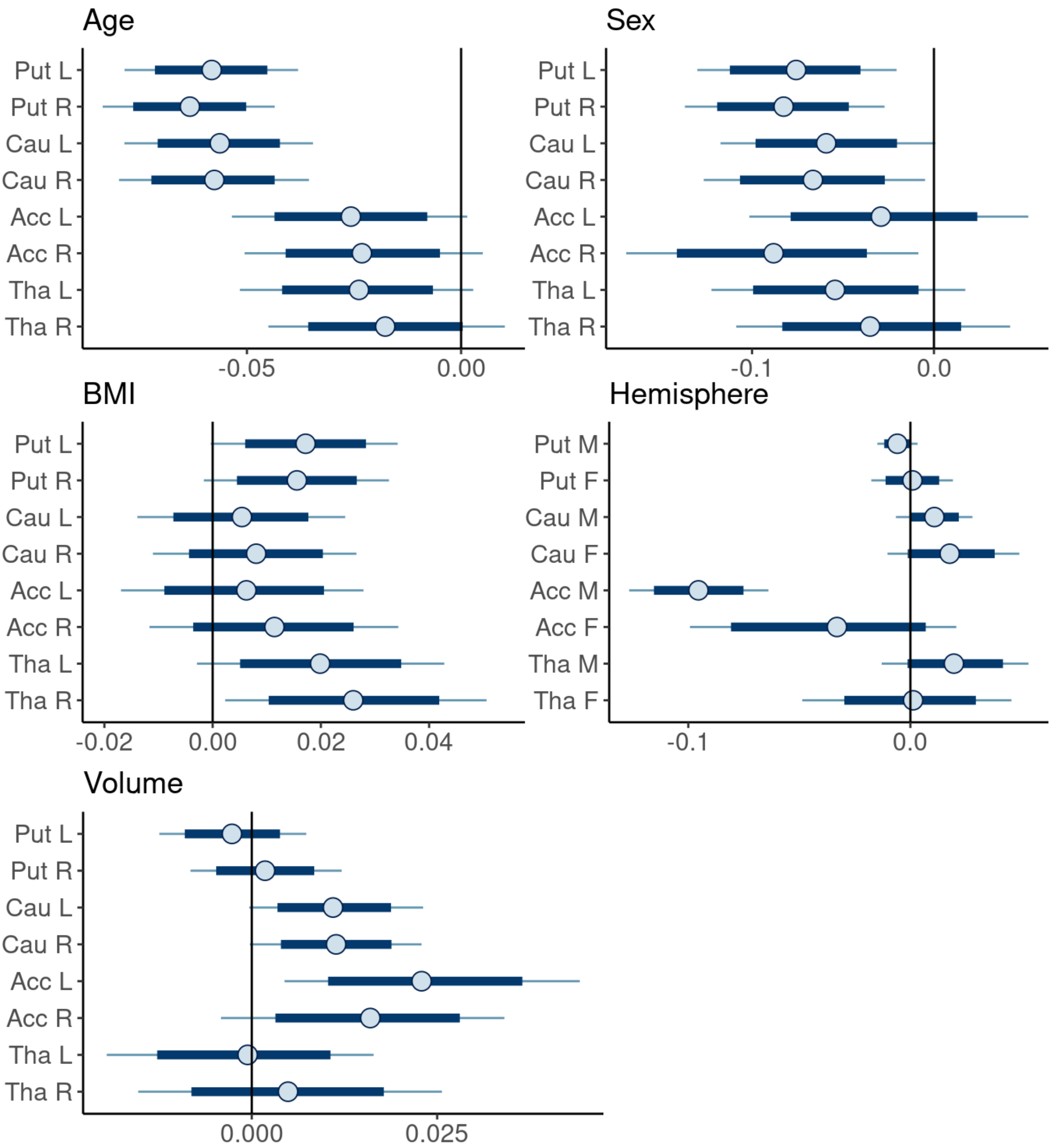


#### Validation of an alternative approach for defining ROIs and reference regions

The results were validated using two different spatial normalization methods, and the age and sex effects were also replicated in an independent sample (n=135).

The figure below shows the ratios of the area under curve (AUC) estimates between the two spatial normalization methods in the reference area cerebellum for the subjects that MRI was available (n=189). According to the figure, the two spatial normalization methods produce highly similar $BP_{ND}$ estimates to the reference region, as most of the ratio observations are close to 1 (i.e., the AUCs are highly similar for both methods) and as +- 3% from the peak includes approximately 99% of the estimates.

```
y = auc_noNA$auc_pet / auc_noNA$auc_mri
mu = mean(y)
sigma = sd(y)
dy = dnorm(y,mu,sigma)

plot(y, dy,
     xlab = "y= AUC Secondary: AUC Primary",
     ylab = "N(y,μ,σ)",
     cex = 1,
     cex.lab = 1,
     cex.axis = 1)
lines(c(mu,mu),c(0,max(dy)))
```

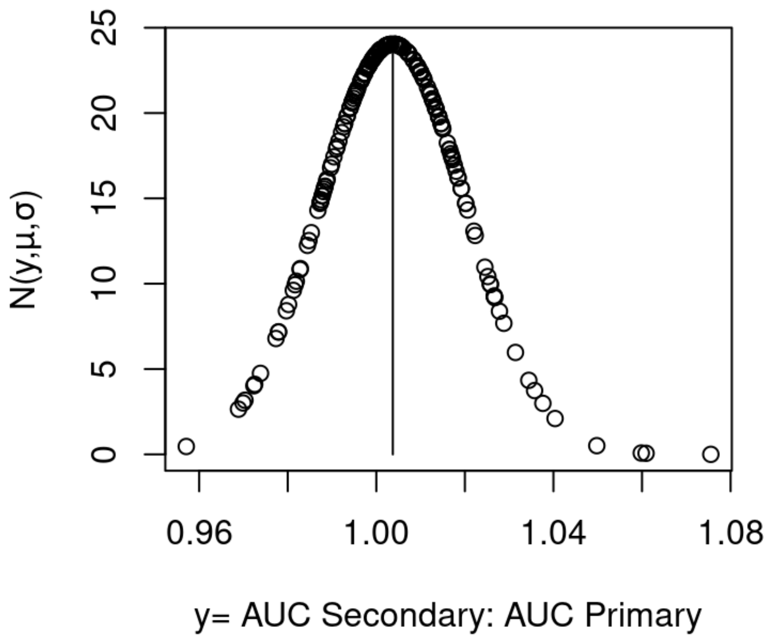


Next, the scatter plots show the high correlation of the two different spatial normalization methods. On the x-axis, there are the $BP_{ND}$ estimates (original scale) acquired from the primary FreeSurfer based normalization method utilizing MR scans. This was the normalization method used in the primary model. On the y-axis, there are the $BP_{ND}$ estimates (original scale) acquired with the alternative method, SPM's "old normalize" tool.

```
ggplot(MRIPET, aes(x=bpMRI, y=bpPET))+
  geom_point(pch=21)+
  geom_smooth(method='lm')+
  geom_abline(intercept=0, slope=1)+
  scale_x_continuous(labels=scales::number_format(accuracy=.1))+
  facet_wrap(~roihemi_f,
             labeller= labeller(roihemi_f = roihemi.labs),
             nrow = 2,
             scales = 'free') +
  theme_classic()+
  theme(strip.background = element_blank())+
  theme(text=element_text(size=12))+
  theme(axis.text=element_text(size=12))+
  theme(strip.text.x=element_text(size=12))+
  labs(title= "", y=expression(paste(BP[ND]," Alternative Method")), x=expression(
paste(BP[ND]," Primary Method"))) +
  theme(aspect.ratio=1)+
  theme(plot.margin=unit(c(0,0,0,0), "cm"))
```

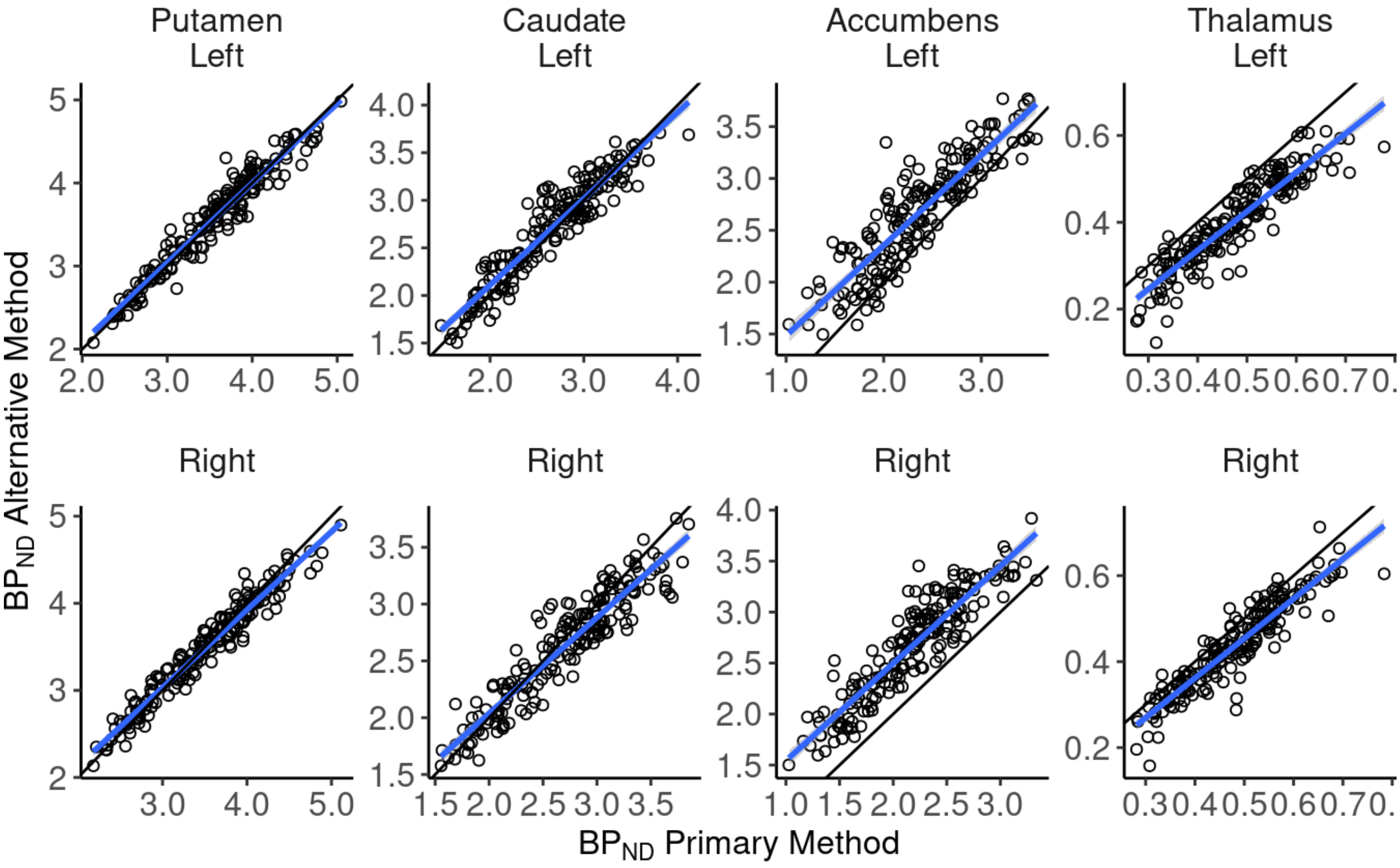


Finally, we tested the replicability of the age and sex effects in the secondary sample (n= 135).

#### Model Settings: Secondary Model

Initially, we executed the secondary modeling with the same parameters and priors as in the primary analysis. However, the model produced 54 divergent transitions. Hence, we modified the parameters to improve the model fitness.

```
rstan_options(auto_write = TRUE)
ITER <- 4000
WARMUP <- 1000
NUM_CORES <- 5
CHAINS <- 5
AD <- 0.999 # modified from 0.99
MAX_TREEDEPTH <- 20

custom_prior <- c(
  set_prior("normal(0,0.5)", class = "b"), # modified from normal(0,1)
  set_prior("normal(0,0.5)", class = "sd") # modified from normal(0,1)
)
```

#### Model: Secondary Model

```
form.atlas.mod <- bf(
  log_bp ~ (1 | subject) + (1 | gr(scanner, id = "scanner")) + (1 | gr(scanner:roi
, id = "scannerroi")) + (1 + age_z + sex | gr(roi, id = "roi")) + age_z + sex,
  sigma ~ (1 | gr(scanner, id = "scanner")) + (1 | gr(roi, id = "roi")) + (1 | gr(
scanner:roi, id = "scannerroi"))
)

#fit.atlas.mod <- brm(
#  formula = form.atlas.mod,
#  data = data2,
#  prior = custom_prior,
#  cores = NUM_CORES,
#  chains = CHAINS,
#  iter = ITER,
#  warmup = WARMUP,
#  control = list(adapt_delta = AD, max_treedepth = MAX_TREEDEPTH)
#)
```

#### Results: Secondary Model

The figure below shows the global effects of age (standardized), sex (male-female) on striatal and thalamic $D_2R$ binding (logarithmic scale) in the subset where the $BP_{ND}$ estimates were produced using the alternative spatial normalization method. The figure shows medians (circles), 80% (thick line) and 90% (thin line) posterior uncertainty intervals of the regression coefficients on a logarithmic scale.

```
ggarrange(plotAatlas, plotBatlas, ncol= 2)
```

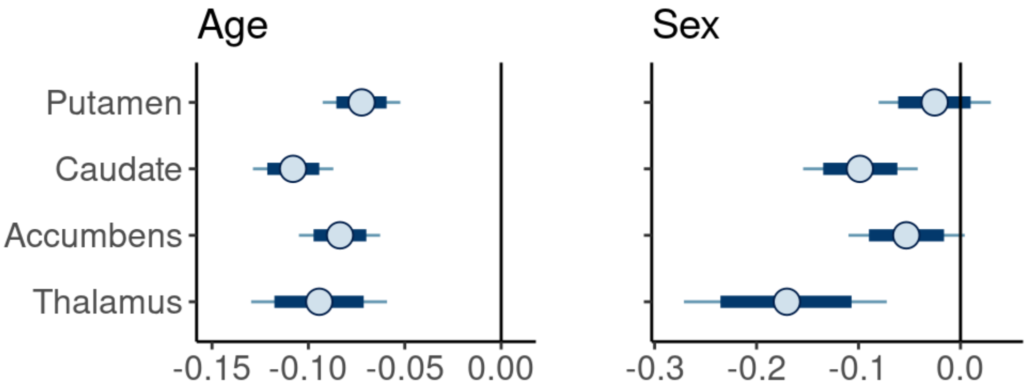

### Between-ROI correlation of $BP_{ND}$ estimates

The figure below shows Pearson correlation coefficients of $BP_{ND}$ estimates between the ROIs. The figure shows that the $BP_{ND}$ estimates are higly correlative between the ROIs.

```
corr <-round(cor(colnames_corr), 2)
p.mat <- cor_pmat(colnames_corr)

ggcorrplot(corr, method = "square",
           hc.order= TRUE,
           type= "lower",
           outline.color= "white",
           lab= TRUE,
           p.mat= p.mat,
           insig="blank",
           ggtheme = ggplot2::theme_classic(),
           show.diag = TRUE,
           lab_size = 4,
           tl.cex = 12)+
  scale_fill_gradient2(limit=c(0.5,1), low= "white", high= "red", midpoint = 0.6,
name = "")+
  theme(text=element_text(size=12))+
  theme(axis.text=element_text(size=12))+
  theme(strip.text=element_text(size=12))+
  theme(text=element_text(size=12))
```

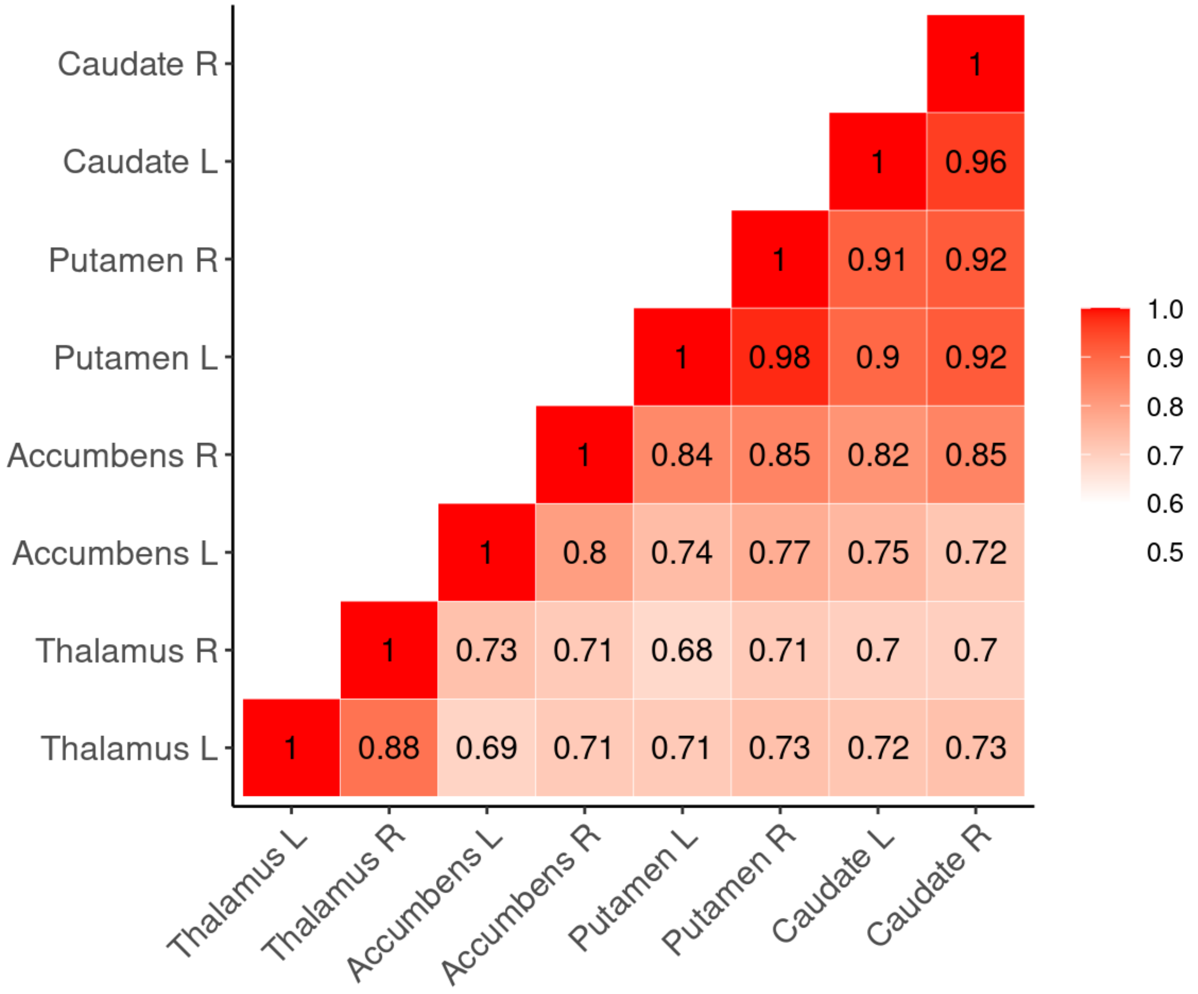

### Conclusions

This RMarkdown document presented the statistical analysis and visualization in our study Age and sex dependent variability of type 2 dopamine receptors in the human brain - a large-scale PET cohort. In the study we concluded that the $D_2R$ availability is dependent on subject characteristics, such as age and sex, and that those differences may contribute to the vulnerability to neuropsychiatric disorders. We also validated an alternative method to normalize PET data if MR image is not available.
